# Neurocomputational mechanisms of affected beliefs

**DOI:** 10.1101/2021.10.26.465922

**Authors:** Laura Müller-Pinzler, Nora Czekalla, Annalina V Mayer, Alexander Schröder, David S Stolz, Frieder M Paulus, Sören Krach

**Affiliations:** Department of Psychiatry and Psychotherapy, Social Neuroscience Lab, University of Lübeck, Ratzeburger Allee 160, D-23538 Lübeck, Germany

**Keywords:** belief formation, self-efficacy, affect, computational modeling, fMRI, pupillometry, anterior insula, embarrassment, pride

## Abstract

The feedback people receive on their behavior shapes the process of belief formation and self-efficacy in mastering a given task. However, the neural and computational mechanisms of how the subjective value of these beliefs, and the corresponding affect, bias the learning process remain unclear. We investigated these mechanisms during the learning of self-efficacy beliefs using fMRI, pupillometry, and computational modeling, and by analyzing individual differences in affective experience. Biases in the formation of self-efficacy beliefs were associated with affect, pupil dilation, and neural activity within the anterior insula, amygdala, ventral tegmental area/ substantia nigra, and mPFC. Specifically, neural and pupil responses map the valence of the prediction errors in correspondence with individuals’ experienced affect and learning bias during belief formation. Together with the functional connectivity dynamics of the anterior insula within this network, our results hint at neural and computational mechanisms that integrate affect into the process of belief formation.

## Introduction

Self-efficacy expectation can be defined as a person’s subjective conviction that he/she can overcome challenging situations through his/her own actions^1^. To successfully perform goal-directed actions, humans must learn from incoming information, thereby forming beliefs about the world and about themselves enmeshed in this world. According to economic theory, learning should result in accurate beliefs that represent an internal model of the world that is suitable to inform decision making. Novel theoretical frameworks, however, emphasize that besides the instrumentality (i.e. accuracy) of beliefs, they may also carry intrinsic value in and of themselves^2^, thus shaping the learning process and how people ultimately arrive at their beliefs^3^. In this regard, affective states, such as happiness about one’s own good health prognosis, represent intrinsic values that individuals are inclined to optimize during belief formation^2, 4^. To demonstrate this entanglement of affect and belief formation, we applied a learning task that induces affective reactions during the process of forming conceptually novel beliefs about one’s abilities to master a task^5, 6^. Specifically, we focused on the primary affective states elicited by self-related beliefs – the self-conscious emotions of embarrassment and pride – and their impact on the beliefs. By exerting experimental control over failures and successes during the process of belief formation, we were able to assess how experienced affect relates to computational mechanisms of belief formation and the underlying activity of neural systems, thus explaining the shifts in preferences for information of positive or negative valence during learning.

Affective states are considered to guide cognitive processing, representing embodied and experiential information about the positive or negative value of what people encounter^7, 8^. It is proposed that this internal affective information is integrated with external information to shape beliefs that rather than being objective, are motivated and biased by subjective feelings about the beliefs themselves, leading to a recursive influence of beliefs and affective states on each other^2, 9^. Previous studies supported aspects of Bromberg-Martin & Sharot’s framework by demonstrating that internal beliefs and external feedback can elicit emotions like happiness, pride, or embarrassment^10–13^. Affective states also alter decision making^12, 14, 15^ and cognitive processes like situational judgments or learning styles^7^. Social anxiety, low self-esteem, or depression, which are likely associated with more negative affective reactions to self-related beliefs, have also been found to bias belief formation^5, 16–18^. These findings provide support for the overall rationale of the formation of “affected beliefs”, that is, the notion that beliefs are fundamentally shaped by affective experiences. However, the question remains open of which neurophysiological mechanisms can explain how emotions elicited during learning are associated with biases in belief formation.

Neuroscientific studies provided initial evidence that common brain areas map the value of stimuli, actions, and their motivational relevance during social and non-social learning and decision making^19, 20^. Prediction errors, that is, the mismatch of prior expectation and a situation’s outcome, are minimized by updating beliefs during learning. These are generally processed in the dopaminergically innervated ventral striatum, but also in the orbitofrontal cortex or the amygdala during learning^20–23^. However, more recent findings suggest that there are distinct and unique neural computations which potentially reflect the impact of the prominent motivational and emotional processes during belief formation. For example, studies have shown that distinct value-related neural processes in subregions of the anterior cingulate cortex (ACC) are recruited depending on whether information about oneself or another agent is processed^24, 25^. Other findings revealed that activation in the ventral striatum was modulated when the social context changed from a private to a public situation, suggesting that the presence or absence of other people influenced the sensitivity to the reward value of certain decisions^26^. Biases specific to self-related belief updating, which are absent when one is learning about another person^27^, have been associated with differences in the tracking of negative prediction errors^28^. In this regard, the ventromedial prefrontal cortex (vmPFC) shows valence-specific encoding of self-related information, which has been shown to predict an optimism bias in belief updating^27, 29^.

Affective states triggered after personal failures or successes are particularly important when people acquire novel self-concepts^30^ and develop an initial understanding of themselves as being self-efficacious individuals in a novel task environment. Central to the entanglement of affect and such self-efficacy beliefs is the assumption that people are highly motivated to perform well and maintain or even construct a positively shaped self-image^31, 32^. Within this process, performance feedback elicits self-conscious emotions, such as pride in the case of success^12, 33, 34^, but also embarrassment if one fails to achieve the expected outcome^13, 33, 35^. It has been demonstrated that these emotions are not only a consequence of the situation but also directly affect behavior. Pride experiences function as a motivator to persevere ^34^. In contrast, embarrassment experiences rather lead individuals to stop their current behavior, withdraw, and appease others^36, 37^. For the process of belief formation, it has been argued that specifically the dorsomedial frontal cortex (dMFC), the ventral and dorsal anterior insula (vAI/ dAI), and the amygdala, brain areas involved in action monitoring as well as emotional processing, integrate affective states with outcome information^38^. Therefore, the anterior insula (AI) has been regarded, among other brain regions, as an integrative hub for motivated cognition and emotional behavior^38, 39^. Similarly, dopaminergic midbrain nuclei in the ventral tegmental area and substantia nigra (VTA/ SN) are associated with attention processes, and at the same time, with events (i.e. reward cues) that are of motivational significance specifically during learning^40, 41^.

While current frameworks support the idea that intrinsic outcomes such as affective states impact the process of belief formation^2, 4^, studies on this issue have not yet probed this framework as a whole. We aim to bridge this gap by showing how emotional states relate to biases in self-related beliefs, that is, the formation of self-efficacy, and how they shift preferences for information of positive or negative valence. For this purpose, we tested the effects of individual differences in affective reactions to the task and to learning behavior. Affective states were evoked during the learning of self-efficacy beliefs in a conceptually novel task environment. Using trial-by-trial updates of performance expectations, we computed prediction error learning rates by fitting computational learning models, revealing valence-specific learning biases. As predicted by current frameworks, individual differences in the experience of the emotions embarrassment and pride were distinctly related to biases in the learning of self-related beliefs. Biased belief updating and affect were jointly related to the neural processing of valence-specific prediction errors in the AI, amygdala, VTA/ SN and mPFC as well as pupillary reactivity in favor of the preferentially used information to update the belief. Increases in valence-specific functional connectivity of the dAI with the amygdala, VTA/ SN and mPFC support the notion of an integrative mechanism of affective and attentional processes within the dAI. These findings provide insights into brain networks involved in computational biases shaped by emotional experiences, and coherently support current theoretical frameworks integrating affective experiences in the process of belief formation.

## Results

### Experimental design

About half of the participants completed the task in the MRI while eye-tracking data were additionally assessed during scanning, and the other half completed the task as a behavioral study (see Methods section for details). In our self-efficacy learning experiment, the “learning of own performance” (LOOP) task^5^, participants were repeatedly confronted with manipulated feedback on their own and another person’s cognitive estimation ability. In different domains (e.g., estimating the weight of animals) participants were led to form novel beliefs about their performance-related self-efficacy. Each participant, together with a confederate, took part in the “cognitive estimation” experiment in our neuroimaging lab. The participant performed the task in the MRI scanner while the confederate (introduced as another participant) allegedly simultaneously performed the task in an adjacent room. During the task, participants were asked to estimate specific characteristics of different properties (e.g., heights of buildings or weights of animals). After each trial, they received manipulated performance feedback for their last estimation (see Figure 1a). During the entire experiment, participants took turns in performing the estimation task themselves (“Self” condition) or observing the other participant performing the task (“Other” condition). Before each trial, participants were requested to rate either their own or the other person’s expected performance for the upcoming trial, enabling us to examine the process of self-and other-related belief formation. The design of the LOOP task provided a High Ability and a Low Ability condition, resulting in overall four feedback conditions: Agent condition (Self vs. Other) x Ability condition (High Ability vs. Low Ability; see Figure 1b and the Methods section for a detailed description of the task). In previous studies, we showed that over time, participants adapted their expected performance ratings according to the feedback, allowing for an assessment of valence-specific self- and other-related learning processes^5, 6^.

**Figure 1.**
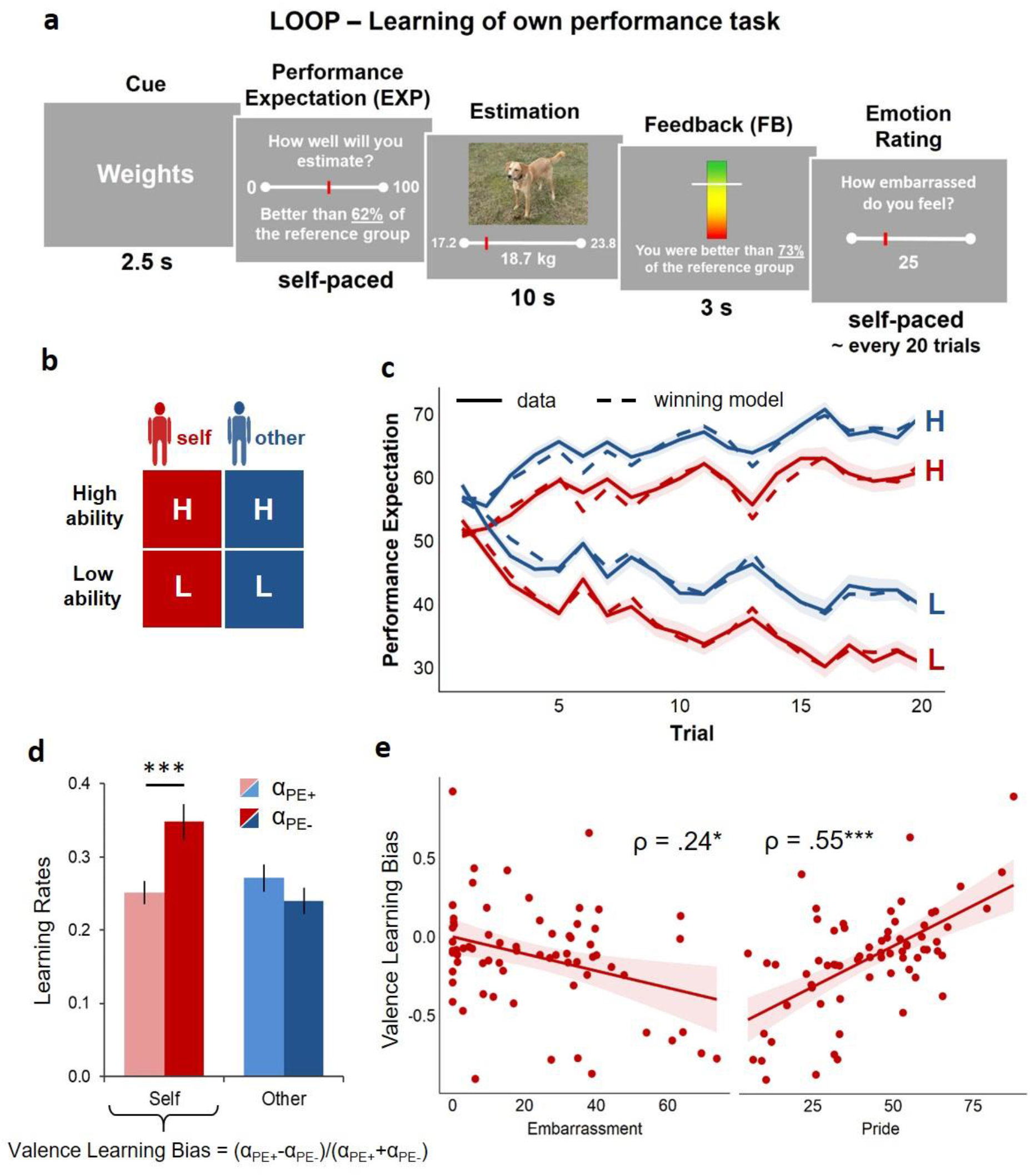
Trial sequence and timing, experimental conditions, modeling of learning behavior, learning rates and their association with self-conscious emotions. **a)** A cue at the beginning of each trial indicated the following estimation category and the agent whose turn it was. After providing their performance expectation ratings (EXP), participants were asked an estimation question, followed by the corresponding performance feedback. After approximately every 20 trials, participants were asked to rate their current emotional state (pride, embarrassment, happiness, stress/ arousal). **b)** The LOOP task contained two experimental factors, Ability level (High Ability vs. Low Ability) and Agent (Self vs. Other), resulting in four feedback learning conditions that can be distinguished by different estimation question types (e.g. estimation of weights). **c)** Predicted and actual performance expectation ratings across time. The behavioral data indicate that participants adapted their performance expectation ratings (solid lines) to the provided feedback, thus learning about the allegedly distinct performance levels. The winning valence-specific learning model captured the participants’ behavior, as indicated by a close match of actual performance expectations with the predicted data (dashed lines). Shaded areas represent the standard errors for the actual performance expectations. **d)** Learning rates derived from the winning Valence Model indicate that there was a bias towards increased updating in response to negative prediction errors (α_PE-_) in contrast to positive prediction errors (α_PE+_) for self-related learning. Bars represent mean learning rates, error bars depict +/- one standard error; *** = *p* < .001, indicates a significant negativity bias during self-related learning. **e)** Correlation plots and Spearman correlations of self-related Valence Learning Bias and embarrassment as well as pride experience during the experiment. * = p < .05, *** = p < .001.

### Selection of computational models for self-related learning

Following a model free behavioral analysis (**Supplementary Results**), we modeled the participants’ behavior by means of learning rates. Therefore, we used the trial-specific expectation ratings for Self and Other to model prediction error (PE) update learning and to assess differences between updating behavior for information of positive vs. negative valence. Our model space contained different models, allowing us to assess the importance of valence-specific learning rates in contrast to unbiased learning between conditions and agents (**Figure S1**). In line with our previous studies, an extended version of the Valence Model, including separate learning rates for positive and negative PEs for Self vs. Other, was the winning model (Model 8; for a more detailed description of this model and the whole model space, see Methods section). This model received the highest sum PSIS-LOO score (approximate leave-one-out cross-validation (LOO) using Pareto smoothed importance sampling (PSIS))^42^ out of all models (for all PSIS-LOO scores see **Supplementary Table S1**). In addition, Bayesian model selection (BMS) resulted in a protected exceedance probability of *pxp>*.999 for this model and a Bayesian Omnibus Risk of *BOR*<.001. Thus, the extended Valence Model was selected for all further analyses of learning parameters, allowing for a comparison of valence-specific learning rates. The time courses of performance expectation ratings predicted by our winning model successfully captured trial-by-trial changes in the actual expectations due to PE updates within each of the ability conditions at the individual subject level (*R^2^*=0.46±0.28 [*M*±*SD]*), supporting the validity of the model in describing the subjects’ learning behavior. In addition to revealing PE valence-specific learning, which could not be directly assessed via model-free behavioral analyses, posterior predictive checks also confirmed that the winning model captured the core effects in our model-free analysis (see **Supplementary Results** and Figure 1c).

### Replication of the negativity bias for self-related learning

Participants showed higher learning rates when learning about themselves than when learning about another person (main effect of Agent: *F_(1,67)_*=5.77, *p*=.019). There was also a main effect of PE valence [pos| neg] (*F_(1,67)_*=5.22, *p*=.025; indicating the categorical comparison of learning rates for positive vs. negative PEs) and a significant interaction of Agent x PE valence [pos| neg] (*F_(1,67)_*=21.47, *p*<.001), which replicates earlier findings of a bias towards more negative updating when learning about one’s own performance (*t_(68)_*=-4.85, *p*<.001, *M*α_PE+Self_=0.25, *SD*=0.13; *M*α_PE-Self_=0.35, *SD*=0.20)^5^. Learning about the other person’s performance did not reveal a significant bias towards more negative updating (*t_(68)_*=1.53, *p*=.128; *M*α_PE+Other_=0.27, *SD*=0.16; *M*α_PE-Other_=0.24, *SD*=0.15; see Figure 1d). There was no significant main effect or interaction for Group (*p*>.097).

### Associations of self-related learning with self-conscious emotions

To quantify associations between learning behavior and affect, individual differences in the overall experience of embarrassment and pride during the task were used as between-subject measures. Embarrassment and pride ratings were only weakly correlated (*ρ_(68)_*=-.10, *p=*.436), indicating that the experience of embarrassment and pride during the task represent two rather independent affective components with respect to the self-related feedback. The bias in self-related learning (Valence Learning Bias=(α_Self/PE+_ - α_Self/PE-_)/(α_Self/PE+_ + α_Self/PE-_))^5, 43, 44^ was negatively linked to the reported experience of embarrassment during the task (*ρ_(68)_*=-.24, *p=*.043; fMRI subsample: *ρ_(38)_*=-.44, *p=*.005), that is, more negative updating behavior was associated with increased embarrassment ratings. In contrast, the Valence Learning Bias was positively linked to the emotion of pride *(ρ_(68)_*=.55, *p<*.001; fMRI subsample: *ρ_(38)_*=.47, *p=*.002). A regression predicting the Valence Learning Bias with both affect ratings simultaneously revealed independent effects of pride (*β*=0.56 *t_(66)_*=5.81, *p<*.001) and embarrassment (*β*=-0.22, *t_(66)_*=-2.30, *p=*.025; *R^2^*=.41, *F_(1,66)_*=22.90, *p*<.001). This indicates that the experience of self-conscious emotions during successful and unsuccessful performances are tied to the way in which people updated their self-efficacy beliefs (see Figure 1e). Furthermore, the way in which participants processed the performance feedback in order to update their self-related ability beliefs was associated with their self-esteem. Specifically, participants with higher self-esteem showed more positive updating, *ρ_(68)_*=.33, *p=*.006 (fMRI subsample: *ρ_(38)_*=.35, *p=*.030), which strengthens the assumption that prior beliefs about the self have a direct impact on how individuals learn novel information about new abilities^5, 45^.

### Pupil dilation slopes are associated with surprise and valence of prediction errors, in line with a negative learning bias

Previous research has successfully linked changes in pupil diameter to surprise, PEs and learning^46, 47^ as well as emotional experiences and arousal^13, 48^. To corroborate our assumption that changes in pupil diameter, as indicated by the slope of the change in pupil size during self-related feedback presentation, reflect increased arousal or attention in association with greater PEs, we regressed trial-by-trial variability in the pupil slope on PE surprise (continuous effect of unsigned PEs) and PE valence [neg*↗* pos] (continuous effect of signed PEs; see Figure 2a)^49^. The linear mixed model revealed a significant positive effect for PE surprise (*t_(1406)_*=2.20, *p*=.028) and a significant negative effect for PE valence [neg↗ pos] (*t_(31.2)_*=-2.50, *p*=.018; see Figure 2b). First, we observed an effect of PE surprise, insofar as the more surprising the feedback was with respect to trial-by-trial prior expectations, the more pupil dilation increased. Second, the results indicate that pupil dilation was greater with decreasing PE values, thus linking negative PEs, rather than positive PEs, to greater dilation. Potentially, these findings indicate increased arousal and attention towards negative PEs, in line with the negativity bias that we found in learning rates.

**Figure 2.**
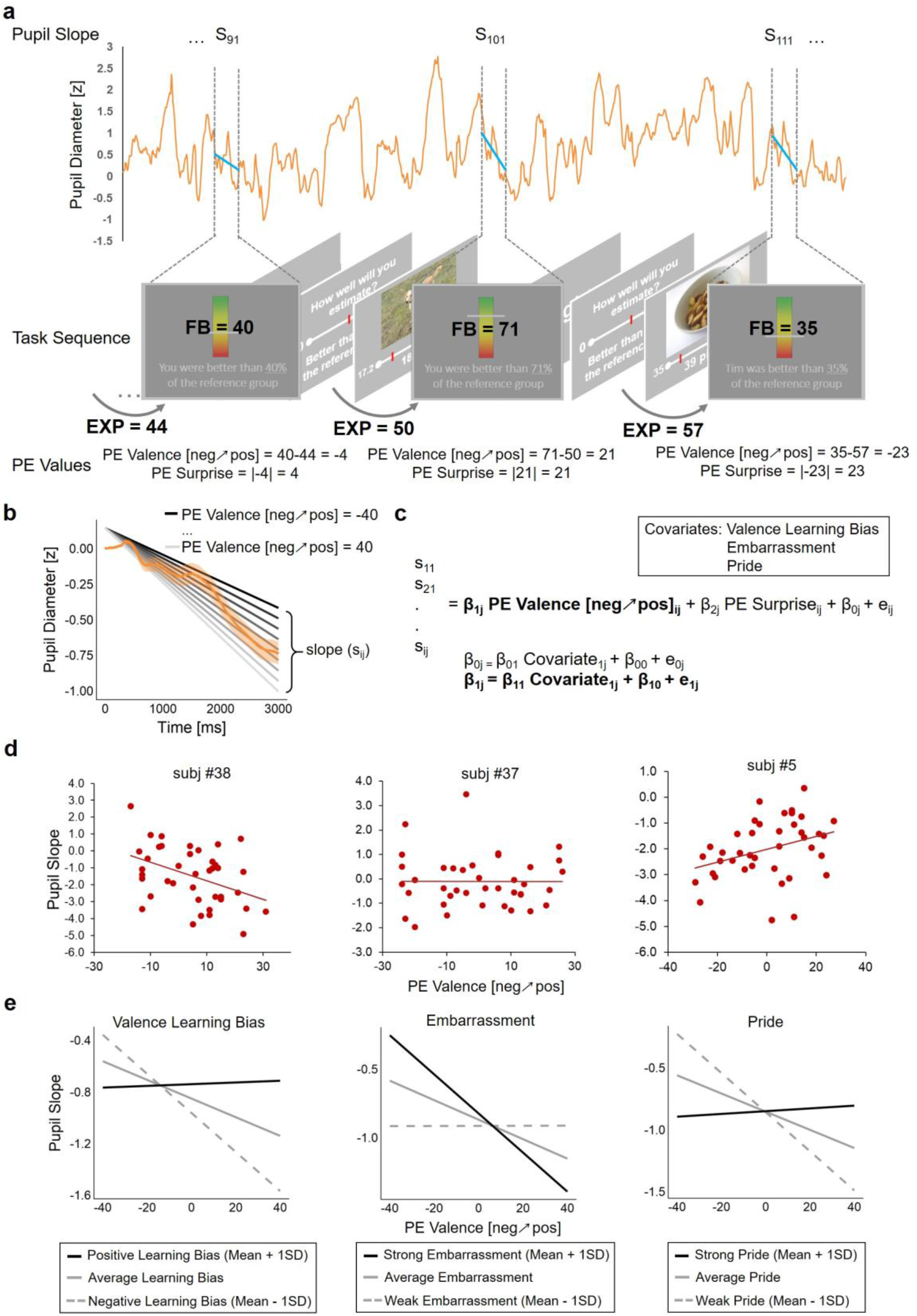
Association of pupil slopes with prediction error (PE) valence and individual pupil response differences explained by differences in Valence Learning Bias, embarrassment and pride experience. **a)** Example of pupil diameter trace over three trials for one subject (orange line) and trial-specific fitted linear slopes (blue lines) for the feedback phase of each trial. PE values are calculated with the participant’s current performance expectation (EXP) and the following feedback value (FB), and PE valence [neg↗ pos] represents the signed PE while PE surprise represents the unsigned PE. **b)** More negative PEs are associated with greater pupil slopes compared to positive PEs. The average pupil diameter trace during feedback is depicted in orange, the shaded area represents +/- one standard error. Pupil slopes for the different levels of PEs (from black = negative to grey = positive) were predicted by the multi-level model containing PE valence [neg↗ pos] and PE surprise as predictors, as described in the Methods section and depicted in **c**. **c)** Description of the multi-level model assessing the association of PE valence [neg↗ pos] and PE surprise with within-subject trial-by-trial pupil slopes and the impact of Valence Learning Bias, embarrassment and pride experience as between-subject second-level covariates explaining differences in the associations on the within-subject level (cross-level interaction; indicated in bold). **d)** Three exemplary scatter plots show the association of pupil slopes with PE valence [neg↗ pos] and illustrate the variance between subjects. **e)** Illustration of the impact of the three between-subject covariates, Valence Learning Bias (left), embarrassment (middle) and pride experience (right) explaining differences in the associations of PE valence [neg↗ pos] and pupil slope. The plots show the data as predicted by the multi-level models for the mean covariate (grey line) and the mean covariate +/- 1 standard deviation (SD; black line and gray dashed line).

### Pupil dilation response to prediction error valence is associated with affect and learning bias

It has been suggested that pupil dilation reflects differences not only between stimuli but similarly between individual biases during decision making (see Figure 2d for examples of individual differences (see Figure 2d for examples of individual differences)^50^. We thus introduced individual differences in learning and self-conscious emotions as between-subject covariates into the linear mixed models assessing trial-by-trial pupil slopes (see Figure 2c). These analyses demonstrated that individuals who experienced more embarrassment showed stronger pupil dilations for negative compared to positive PEs, while pupil slopes did not differ between positive and negative PEs in individuals with lower embarrassment (significant interaction of embarrassment and PE valence [neg↗ pos]: *t_(31.8)_*=-2.57, *p*=.015; no main effect for embarrassment: *t_(34)_*=-0.42, *p*=.680; **see** Figure 2e). These effects were reversed when pride ratings, instead of embarrassment ratings, were included in the model (interaction pride and PE valence [neg↗ pos] *t_(32.8)_*=3.14, *p*=.004; main effect of pride: *t_(34.1)_*=0.04, *p*=.971). The Valence Learning Bias modulated the relationship between PE valence [neg↗ pos] and pupil slopes in the same way (interaction Valence Learning Bias and PE valence [neg↗ pos] *t_(31.3)_*=2.96, *p*=.006; main effect of Valence Learning Bias: *t_(34.3)_*=1.05, *p*=.300), indicating that participants with a more negative Valence Learning Bias had greater pupil dilation for negative PEs, whereas participants with no bias or a positive bias showed less differentiation in pupil dilation in response to the valence of the PE.

### Common neural activations associated with prediction error surprise and distinct activations for self-related prediction error valence

We assessed the effects of continuous trial-by-trial PE surprise and PE valence [neg↗ pos] as parametric weights to assess neural aspects of learning more specifically (see Figure 3a). Increased PE surprise was associated with greater activation of the mPFC for Self and Other as well as clusters in the bilateral insula/ temporal pole/ frontal orbital gyrus at a more lenient threshold (cluster-wise FWE-corrected with p < .05 at a cluster forming threshold of p < .001; see Figure 3c and **Supplementary Table S4**). There was no significant difference between Self and Other (*p* < .001), suggesting that there may be a common process of error tracking mapped within the same brain regions, independent of the agent.

**Figure 3.**
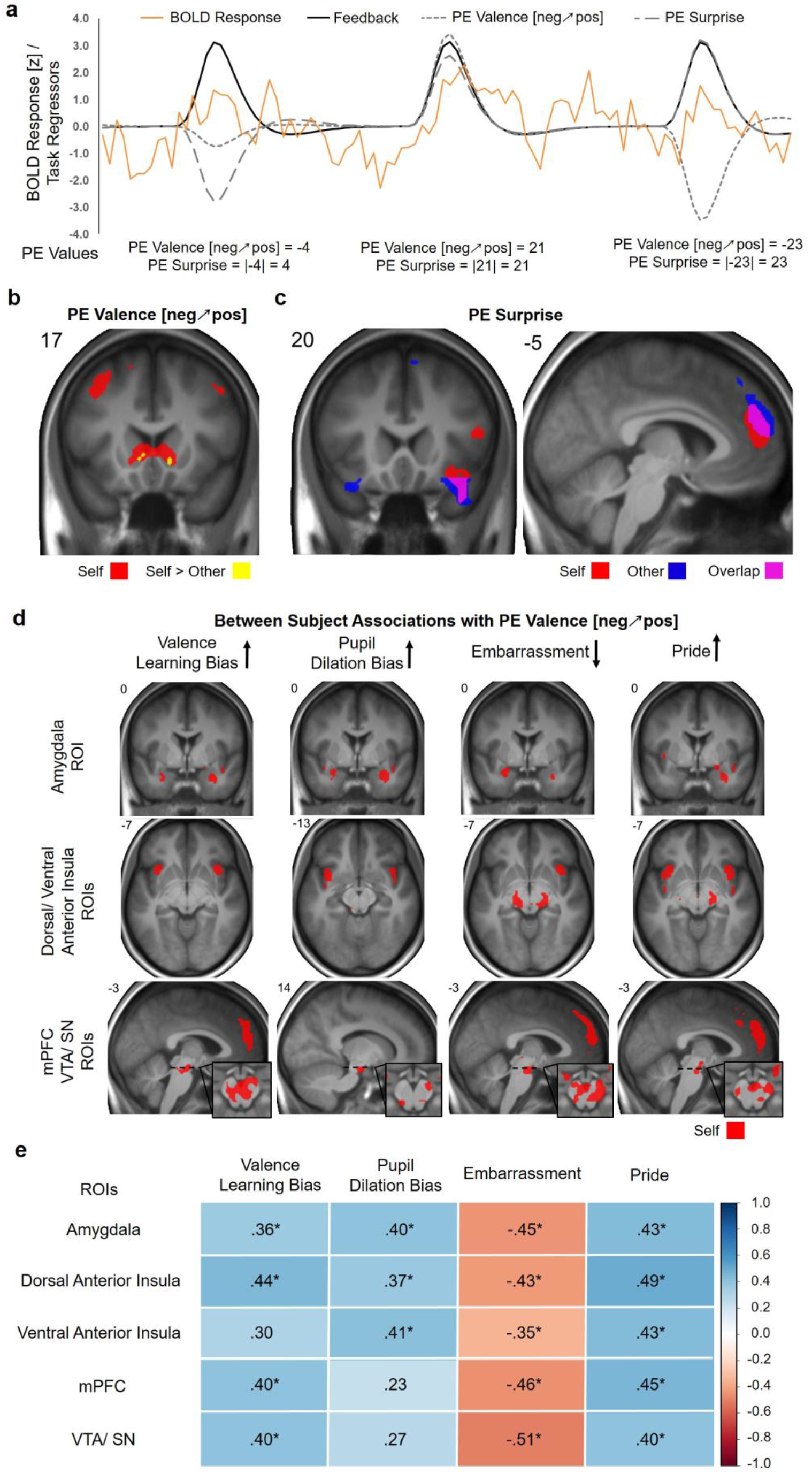
Common neural activations associated with prediction error (PE) surprise, distinct activations for self-related PE valence [neg↗ pos] and individual response differences to PE valence [neg↗ pos] explained by differences in Valence Learning Bias, embarrassment and pride experience, and pupil dilation. **a)** Exemplary BOLD response over three trials for one subject (orange line) and regressors for the feedback phase of each trial (back line; the originally separate regressors for self-and other-related feedback are combined here for display purposes). PE valence [neg↗ pos] (small dashed) and PE surprise (large dashed) are added as parametric modulators in addition to the feedback regressors. PE values are calculated as shown in Figure 2. **b)** PE valence [neg↗ pos] was associated with increased activation of the NAcc/VS, mPFC, bilateral angular gyrus/ superior parietal lobule/ lateral occipital gyrus and precentral gyrus for Self. Activation of the NAcc/VS was stronger for Self than for Other. **c)** PE surprise was associated with activation of the mPFC and the bilateral insula/ temporal pole/ frontal orbital gyrus for Self and Other. **d)** Neural tracking of self-related PE valence [neg↗ pos] in the predefined regions of interest was modulated by between-subject variables. Black arrows indicate the direction in which the covariates are coded in the analyses. Clusters refer to p < .005, uncorrected for display purposes; see **Supplementary Table S7** for FWE-corrected statistics. **e)** Pearson correlations for parameter estimates derived from the whole areas of our predefined ROIs with the Valence Learning Bias, Pupil Dilation Bias, embarrassment and pride are color-coded. * = p < .05, FDR corrected.

The assessment of PE valence [neg↗ pos] revealed a distinct pattern for self- and other-related learning: Self-related PE valence [neg↗ pos] was positively associated with increased activation of the NAcc/VS, mPFC, bilateral angular gyrus/ superior parietal lobule/ lateral occipital gyrus and precentral gyrus, showing a stronger activation for positive vs. negative PEs (Figure 3b and **Supplementary Table S5**). There was no effect for other-related PE valence [neg↗ pos], and a direct comparison of self-vs. other-related PE valence [neg↗ pos] effects revealed stronger associations in the NAcc/VS for Self (right: x, y, z: 12, 17, −4, t_(38)_ = 5.23; k = 2; left: x, y, z: −9, 26, −1, t_(38)_ = 5.77, k = 19), supporting the assumption that the valence of the feedback has a specific value when feedback refers to the self as compared to another person. Although behavioral data and learning rates clearly emphasize greater importance of negative over positive PEs, there were no significant negative associations with PE valence [neg↗ pos] in the neural data (*p* < .001). To test whether the activations associated with self-related PE valence [neg↗ pos] were actually related to PEs and not merely to the feedback value alone, we ran an additional control analysis including parametric weights for trial-by-trial feedback and performance expectation ratings instead of PE values^51^. This model replicated the findings, showing positive associations within the reported brain regions, including the NAcc/ VS, with trial-by-trial self-related feedback values and negative associations with prior expectations as would be expected for PE-related effects (**Supplementary Table S6**).

### Neural activity in response to self-related prediction error valence is associated with affect, learning bias, and pupil dilation

To assess how biases in learning as well as affective experience and pupil dilation were associated with valence-specific PE processing on the single trial level, multiple general linear models (GLMs) were performed. These included the Valence Learning Bias, self-conscious emotions, and a score representing a valence bias for pupil dilation responses to positive vs. negative PEs (Pupil Dilation Bias = PupilSlope_Self/PE+_ - PupilSlope_Self/PE-_) as between-subject covariates for PE valence [neg↗ pos] tracking. Analyses within our predefined regions of interest (ROIs) revealed that the more negative the Valence Learning Bias was, the more neural activity increased in response to negative vs. positive PEs in the bilateral dAI, vAI, amygdala, mPFC, and VTA/ SN (all results are *p*<.05 FWE-corrected within ROIs; see Figure 3d and **Supplementary Table S7**). Overall, higher experience of embarrassment showed similar associations with increased PE tracking for negative vs. positive PEs in the right dAI, bilateral amygdala, and VTA/ SN. Trend effects for embarrassment were found in the left dAI, bilateral vAI, and mPFC. In line with this, lower experience of pride showed the same association in the dAI and vAI, amygdala, VTA/ SN and mPFC. Additional analyses revealed that effects for embarrassment and pride were mainly independent (see **Supplementary Results**). Similarly, the more negative the Pupil Dilation Bias was, the stronger the increase in activation of the dAI and vAI, amygdala and VTA/ SN towards negative vs. positive PEs. Thus, the greater the response of this neural system to PEs with negative valence, the greater was the preference for negative information during learning as well as the negativity of the affective experience. This gained multi-modal support by similar associations of the Valence Learning Bias and affect with the pupil dilation response, which reflects the activity of this underlying neural system. In contrast, participants who showed a greater response of this neural system to positive PEs also had a preference for positive information during learning and reported more positive affect.

### Functional connectivity of the dorsal anterior insula depends on prediction error valence in line with the negativity bias

Functional connectivity dynamics of the left and right dAI were assessed, as these were activated during feedback processing for self- and other-related feedback, independent of Agent and PE valence. Using psychophysiological interaction (PPI) analyses, we tested whether the connectivity of the dAI differed depending on the level of the valence of the PE. We calculated the interaction of the continuous PE valence [neg↗ pos] and the time series extracted from the left and right dAI seed regions separately for Self and Other on the first level. The two agents were contrasted against each other on the second-level GLM, as we were specifically interested in connectivity dynamics that might reflect the differential learning from negative PEs when processing self-relevant information. Contrasting the PPI effects for PE valence [neg↗ pos] between Self and Other demonstrated that during self-related learning, functional connectivity dynamics of the right dAI with the bilateral amygdala, mPFC and VTA/ SN (p<.05, FWE-corrected within ROIs) more strongly aligned with the negativity of the PEs. The left dAI showed a weaker but similar spatial distribution, with significant differences between self- and other-related PE valence [neg↗ pos] for the left amygdala and VTA/ SN (p <. 05, FWE-corrected, see Figure 4a/ b and **Supplementary Table S8**). Thus, those brain regions that preferentially tracked PEs of negative valence in individuals with increased negative affect and learning biases also showed connectivity dynamics with the dAI in a similar direction during self-related learning. Individuals who showed more pronounced differences in functional connectivity, that is, stronger functional connectivity for negative PEs during Self>Other, also showed a more negative Valence Learning Bias, although this pattern was not fully consistent across all ROIs (see Figure 4c).

**Figure 4.**
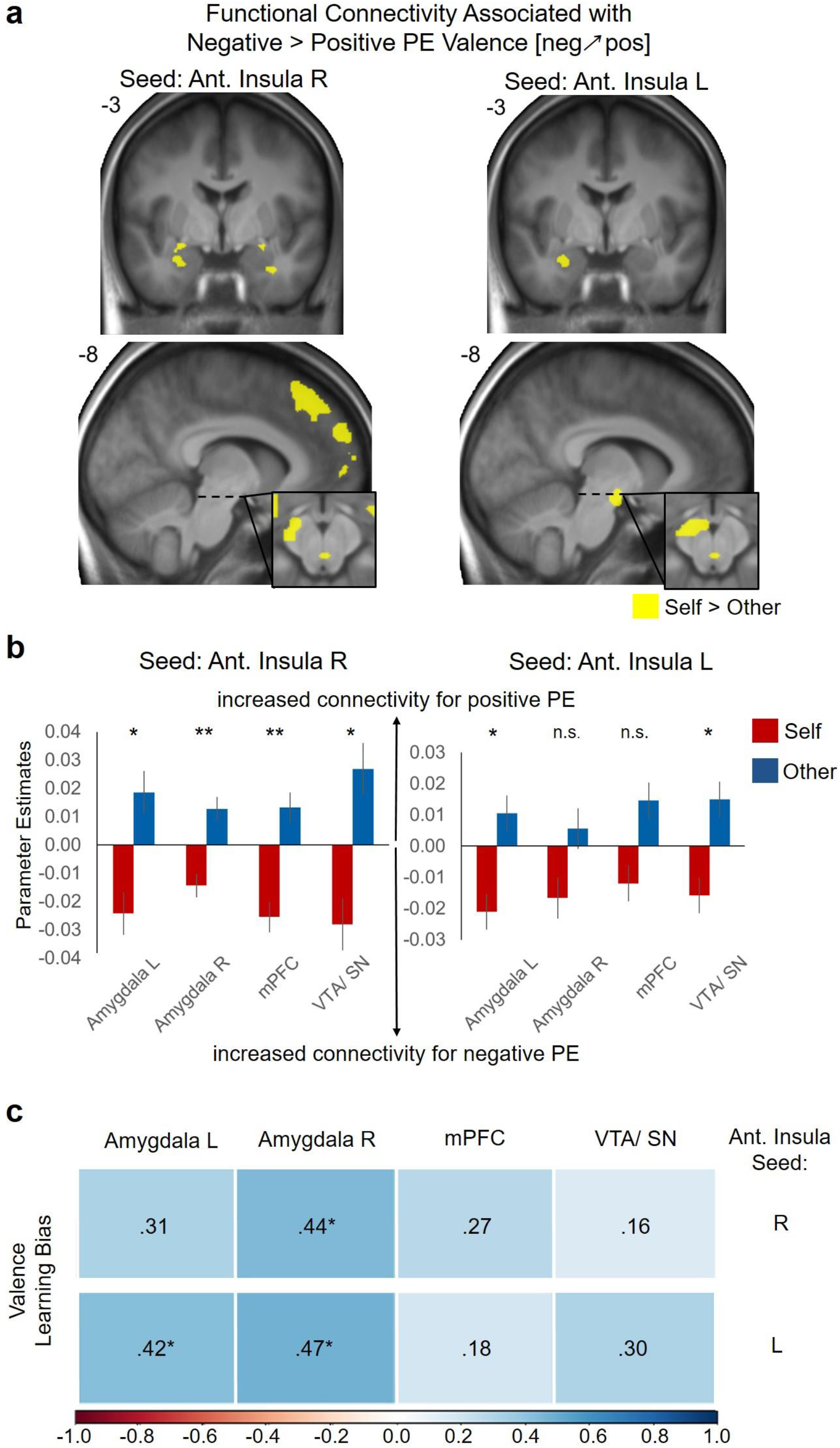
Differences in functional connectivity of the dorsal anterior insula during prediction error (PE) valence [neg↗ pos] tracking for self- and other-related learning and associations with the Valence Learning Bias. **a)** Increased functional connectivity of the dorsal anterior insula for the negative effect of PE Valence [neg↗ pos] for self- vs. other-related learning in the predefined ROIs (amygdala, mPFC, VTA/ SN; p < .005 uncorrected for display purposes). **b)** Functional connectivity dynamics of the dorsal anterior insula plotted separately for self- and other-related learning. For display purposes, parameter estimates are plotted separately for Self and Other and refer to the peak voxels of the contrast Self vs. Other that are reported in **Supplementary Table S8**. * = p < .05, ** = p < .01. **c)** Spearman correlations of the Valence Learning Bias with the functional connectivity dynamics between the dorsal anterior insula (seed region reported on the right side) and the amygdala, mPFC and VTA/ SN associated with PE valence [neg↗ pos] for self- vs. other-related learning are color-coded. * = p < .05, FDR corrected.

## Discussion

Belief formation is essentially biased, and various studies have shown how it is shaped by motivations^3, 32, 52, 53^. The findings of the present study reveal that the affect which people experience during learning is linked to the process of self-related belief formation on the level of neural systems. Our computational modeling results imply that biases in the formation of self-efficacy beliefs in mastering a conceptually novel task are associated with the experience of the self-conscious emotions of embarrassment and pride. Critically, on the level of neural systems, the valence of PEs was associated with biases in self-related learning and negativity of the affective experience. Individual differences in the response preference for negative PEs, as indicated by the pupil dilation response and activation of the AI, amygdala, mPFC, and VTA/ SN, were associated with a more negative learning bias and negative affective experience, hinting at a neurobiological system that integrates affect during learning.

The novel framework on the “value of beliefs” proposed by Bromberg-Martin and Sharot^2^ nicely details how beliefs elicit emotions, while at the same time, these emotions shape how beliefs are updated in a reciprocal relationship. Based on this framework as well as previous research on self-conscious emotions, a negative belief about one’s abilities should elicit stronger embarrassment after failures and reduced pride after successes^13, 33^. According to the present data, the association of the learning bias with the affective experience during learning supports this notion, as individuals who experienced more negative affect (embarrassment) and less positive affect (pride) when receiving self-efficacy feedback were also inclined to update their beliefs in a more negative way. At the same time, negative emotions guide the information processing at various stages, including perception, attention, and decision-making, as discussed in the context of “motivated cognition”^4^. This reciprocal relationship finally results in biased belief formation and in beliefs that are both drivers of affect and influenced by emotional responses to incoming information. Embarrassment is a particularly relevant example illustrating this recursive relationship: The fear of failure, as is often discussed in the context of social anxiety (disorder)^5, 13, 16, 54^, leads to shifts in expectations and attention (threat monitoring) towards negative information. At best, this results in reparative behavior and performance improvement^55, 56^, and at worst, it leads to a vicious cycle of fear and pathologically increasing negative beliefs about the self^57^. This is reflected in the present findings, when individuals who experienced more intense embarrassment ended up with lower self-efficacy beliefs.

Emotions shape learning processes in different ways. First, emotions can influence how information is processed in the brain by adaptively shifting attention towards salient aspects of the situation^58, 59^. Second, emotions entail arousal, which intensifies internal rehearsal and evaluations, leading to increased learning^7, 8, 58^, although these processes often interact and are intricately related^4^. The increased pupil dilation in response to negative PEs in our study is in line with both increased salience of and attentional shifts towards negative PEs^46, 47^ or increased arousal elicited by negative PEs^13, 48^. In this regard, we believe that the stronger impact of positive or negative information on pupil responses and brain reactivity maps arousal and affect according to the valence of individual learning biases and affective experiences.

Specifically, the anterior insula (AI) has been suggested to function as an integrative hub for motivated cognition and emotional behavior^38, 39^. While ventral aspects of the AI are associated with affective processing, emotions, and physiological arousal^39, 60–62^, dorsal aspects of the AI are strongly associated with the detection of salient events, allocation of attentional resources, executive working memory^63, 64^ and also (absolute) PEs and uncertainty during learning^65–67^. These findings suggest that the functions of the AI provide a physiological basis for how emotions are translated into biased, motivated, or affected beliefs^38, 39^. A similar role, as a link for the attention-emotion interaction, has also been suggested for the amygdala^38, 59^, which showed similar responses in our task. The functional connectivity dynamics of the dAI, matching the modeled learning rates with a stronger impact of self-related negative PEs, underline the insula’s role as an integrative hub that receives and forwards information affecting information processing in other brain regions.

Tracking of PEs in the dopaminergically innervated VTA/ SN is influenced by motivational factors during learning^40^. The subjective value of self-related information varies significantly between subjects, as indicated by idiosyncratic response patterns of the VTA/ SN to gains or losses^68^. In this line, we believe that the present results reflect individual response tendencies at a very basic level of PE tracking. On higher layers of the computational hierarchy, regions in the ACC and mPFC are also associated with PE tracking and value representation^24, 69, 70^ and have been previously associated with biases in belief updating^27, 29, 71^. Affect and arousal could therefore bias learning on various stages of the computational hierarchy of PE processing, from more basic dopaminergic midbrain responses to more abstract value representations in the neocortex^72^. While the directionality of the effects remains to be determined, the dynamics in the functional connectivity of the dAI suggest a modulatory role in this process. Here, information is forwarded to and/ or integrated from the VTA/ SN and mPFC, the same regions whose response to the valence of PEs was also modulated by differences in learning bias and affective experience. This strengthens the idea that the AI plays a role in shifting responses to negative or positive information in other brain regions (e.g. by shifting attention and by affective tagging) or that it already receives stronger signals in response to PEs of negative or positive valence from midbrain regions and the mPFC.

The tracking of the absolute error, PE surprise^49^, independently of the agent, is in line with the “common currency” assumption^20, 73^ for the positive and negative value of one’s own and others’ performance feedback. The common and valence-independent coding of surprise in the insula and the mPFC might therefore be sufficient to complete the learning task per se. Valence, however, matters when individuals learn about themselves, as indicated by an additional shift in error tracking in the same regions, AI and mPFC, that also track surprise in a valence-independent manner. As a consequence, we observe a robust effect for surprise across individuals, but when people learn in a more negatively biased manner and experience more negative affect, this signal is unbalanced and increases with more negative PEs. This pattern hints at a neurocomputational mechanism of how affect shapes the formation of beliefs, as proposed previously^2^.

Some of the key findings of the present study emerged at the level of individual differences. We observed a wide inter-individual variance in the affective experience during the task and in the Valence Learning Bias, that is, the type of information participants preferentially used to update self-related beliefs. While, on average, we found a negativity bias during self-related belief formation, just under a third of the participants still showed a positive learning bias, underlining the importance of individual factors and the meaningfulness of variability. Studies suggest not only that biases in belief formation differ between tasks^3, 5, 74^ but also that they depend on situational factors like stress^6, 75^. An individual’s ability to adjust his/her current information processing strategy to the context might be adaptive^2^: for example, adaptation to an increased relevance of negative or threat-related information during stress^75^ or coping with a negative self-concept following social stress by means of more self-beneficial belief updating^6^. It might also be adaptive for people who fear negative feedback to pay more attention to failure-related information in order to learn and circumvent potential future failures^52^. However, it is not always straightforward to determine under which conditions a strategy is adaptive or whether the affective experience can ameliorate the individual’s well-being. A maladaptive consequence of biased self-efficacy beliefs becomes apparent in psychiatric disorders such as depression and social anxiety, in which amplified negative updating can lead to persistently distorted self-views and overly negative beliefs about one’s own capabilities in everyday life^16, 17, 76–78^.

Emotions experienced during learning affect computational mechanisms and manifest in distributed neural activity during belief formation. In particular, neural activity of the AI, amygdala, VTA/SN, and mPFC and pupil responses map the valence of PEs in correspondence with the experienced affect and the learning bias that people show during belief formation. The more negative balancing in the functional connectivity dynamics of the dAI during the processing of self-related PEs within this network outline a scaffold for neural and computational mechanisms integrating affect during belief formation. The results of our empirical implementation of the “value of beliefs” framework^2^ have broader implications concerning any context that provides personal evaluations based on behavioral performance. Here, the focus on the affective experience during learning provides a deeper understanding of how feedback manifests in self-related beliefs, which may in turn significantly impact developmental processes and future behavior.

## Materials and Methods

### Participants

The study was approved by the ethics committee of the University of Lübeck (AZ 18-066), was conducted in compliance with the ethical guidelines of the American Psychological Association (APA), and all subjects gave written informed consent. Participants were recruited at the University Campus of Lübeck, were fluent in German, and had normal or corrected-to-normal vision. Two independent samples were recruited, one for the fMRI study and the other for a behavioral study that was added to increase the sample size for the behavioral data. All participants received monetary compensation for their participation in the study. The final sample size for the fMRI sample was 39 participants (26 females, aged 18-28 years; *M*=22.3; *SD*=2.65). We initially recruited 48 participants, but had to exclude six participants who did not believe the cover story of the task and three participants who did not attentively complete the task until the end (e.g. participants reported that they were too tired or the ratings indicated that they stopped responding to the estimation task). The additional behavioral sample consisted of 30 participants (24 females, aged 18-32 years; *M*=23.3; *SD*=3.97). For more details on the sample characteristics, see **Supplementary Table S9**.

### Learning of own performance task

The learning of own performance (LOOP) task enables participants to incrementally learn about their own or another person’s alleged ability in estimating properties. The task was previously introduced and validated in a set of behavioral studies^5^. For the LOOP task, all participants were invited to take part in an experiment on “cognitive estimation” together with a confederate, who was allegedly another participant. In contrast to the fMRI study, for the behavioral study, two participants were invited and tested together instead of introducing a confederate. Participants were informed that they would take turns with the other participant/ confederate, either performing the task themselves (Self) or observing the other person performing (Other). Participants were asked to estimate different properties (e.g. the height of houses or the weight of animals). On a trial-by-trial basis, participants received manipulated performance feedback in two distinct estimation categories for their own estimation performance and for the other person’s estimation performance. Unbeknownst to the participant, one of the two categories was arbitrarily paired with rather positive feedback while the other was paired with rather negative feedback (e.g. “height” of houses = High Ability condition and “weight” of animals = Low Ability condition or vice versa; estimation categories were counterbalanced between Ability conditions and Agent [Self vs. Other] conditions). This resulted in four feedback conditions with 20 trials each (Agent condition [Self vs. Other] x Ability condition [High Ability vs. Low Ability]). Trials of all conditions were intermixed in a fixed order with a maximum of two consecutive trials of the same condition. Performance feedback was provided after every estimation trial, indicating the participant’s own or the other person’s current estimation accuracy as percentiles compared to an alleged reference group of 350 university students who, according to the cover story, had been tested beforehand (e.g. “You are better than 94% of the reference participants.“; see Figure 1a). The feedback was defined by a sequence of fixed PEs with respect to the participants’ “current belief” about their abilities. The “current belief” was calculated as the average of the last five performance expectation ratings per category, which started at 50% before participants actually rated their performance expectation. This procedure led to varying feedback sequences between participants but kept PEs mostly independent of the participants’ performance expectations and ensured a relatively equal distribution of negative and positive PEs across conditions (Self: mean positive PE = 13.6, SD = 1.8 (average number = 20.3); mean negative PE = −12.6, SD = 1.4 (average number = 19.7); Other: mean positive PE = 13.0, SD = 1.3 (average number = 19); mean negative PE = −13.1, SD = 1.1 (average number = 21)). At the beginning of each trial, a cue was presented indicating the estimation category (e.g. “height”) and participants were asked to state their expected performance for this trial on the same percentile scale used for feedback. In order to increase motivation and encourage honest response behavior, participants were informed as part of the cover story that accurate expected performance ratings would be rewarded with up to 6 cents per trial, that is, the better the match between their expected performance rating and their actual feedback percentile, the more money they would receive. Following each performance expectation rating, the estimation question was presented for 10 seconds. During the estimation period, continuous response scales below the pictures determined a range of plausible answers for each question. Participants indicated their responses by navigating a pointer on the response scale with an MRI-compatible computer mouse. Subsequently, feedback was presented for 3 seconds (see Figure 1a). Jittered inter-stimulus intervals were presented following the cue (2-6 * TR (0.992 secs)), estimation (2.5 – 6.5 * TR) and feedback phase (4-8 * TR) for the fMRI task. All stimuli were presented using MATLAB Release 2015b (The MathWorks, Inc.) and the Psychophysics Toolbox^79^. The fMRI task was completed in two separate 20-min sessions with a short break in between.

Before starting the experiment, all participants answered several questions about their self-related beliefs and completed a self-esteem personality questionnaire (Self-Description Questionnaire-III (SDQ-III))^80^. During the LOOP task, participants were also asked to rate their current levels of embarrassment, pride, happiness and stress/ arousal on a continuous scale ranging from not at all (coded as 0) to very strong (coded as 100). Two emotion rating phases followed self-related feedback and two rating phases followed other-related feedback. The two emotion rating phases following self-related feedback were averaged to obtain a rating for the experience of self-conscious affect (embarrassment and pride) during self-related learning. Following the task, participants completed an interview including ratings about self-related beliefs, were debriefed about the cover story, and reimbursed for their time before leaving. The whole procedure took approximately 2 h.

### Behavioral Data Analysis and Modeling

To illustrate the basic effects in our behavioral data, a model-free analysis was performed on the participants’ expected performance ratings for each trial. We conducted a repeated measures ANOVA with the factors Trial (20 Trials) x Ability condition (High Ability vs. Low Ability) x Agent condition (Self vs. Other). In order to control for potential differences between the two samples Group was added as a between-subject factor. All statistical analyses on the behavioral data apart from the modeling procedure were performed using *jamovi* (Version 1.2.27, The jamovi project (2020). Retrieved from https://www.jamovi.org).

Dynamic changes in self-related efficacy beliefs, that is, performance expectation ratings, were then modeled using PE delta-rule update equations (adapted Rescorla-Wagner model)^81^. The model space contained three main models, which varied with regard to their assumptions about biased updating behavior when learning about the self (see **Supplementary Figure S1**). The simplest learning model used one single learning rate for all conditions for each participant, thus not assuming any learning biases (Unity Model). The second model, the Valence Model, included separate learning rates for positive PEs (α_PE+_) vs. negative PEs (α_PE-_) across both ability conditions, thus suggesting that the valence (positive vs. negative) of the PE biases self-related learning. The third model, the Ability Model, contained a separate learning rate for each of the ability conditions, indicating context-specific learning. In addition, learning rates were either estimated separately for Self vs. Other or across Agent conditions. The Valence Model with separate learning rates for Self vs. Other (Model 5), which was the winning model in our previous studies^5, 6^, was further extended by adding a weighting factor that reduced learning rates towards the ends of the feedback scale (percentiles close to 0 % or 100 %), under the assumption that participants would perceive extreme feedback values to be less likely than more average feedback^82^. In the first of these models (Model 7), a linear decrease of the learning rates was assumed, beginning at 50 % and ending at 0 % and 100 %. A weighting factor *w* was fitted for each participant, defining how strongly the linear decrease was present for each individual. Since many of the variables people encounter in everyday life (e.g., many test results) approximately follow a normal distribution with extreme values being less likely, for the second model of this kind (Model 8), we assigned the relative probability density of the normal distribution to each feedback percentile value. Again, a weighting factor *w* was fitted for each individual, indicating how strongly the relative probability density reduced the learning rates for feedback further away from the mean. In contrast to our previous studies in which we implemented the LOOP task with fixed feedback sequences, here, feedback depended on the participants’ current expectations and thus differed between participants and conditions. Reduced learning rates towards the ends of the feedback scale, which may systematically confound learning rates between participants and conditions, were thus accounted for in Models 7 and 8. To test whether the participants’ performance expectation ratings can be better explained in terms of PE learning as compared to stable assumptions in each Ability condition, we included a simple Mean Model, with a mean value for each task condition (Model 9; for more details, see **Supplementary Methods**).

#### Model Fitting

For model fitting, we used the RStan package (Stan Development Team, 2016. RStan: the R interface to Stan. R package version 2.14.1.), which implements Markov chain Monte Carlo (MCMC) sampling algorithms. All of the learning models in the model space were fitted for each participant individually, and posterior parameter distributions were sampled for each participant. A total of 2400 samples were drawn after 1000 burn-in samples (overall 3400 samples; thinned with a factor of 3) in three MCMC chains. We assessed whether MCMC chains converged to the target distributions by inspecting *R̂* values for all model parameters^83^. Effective sample sizes (*n_eff_*) of model parameters, which are estimates of the effective number of independent draws from the posterior distribution, were typically greater than 1500 (for most parameters and subjects). Posterior distributions for all parameters for each of the participants were summarized by their mean as the central tendency, resulting in a single parameter value per participant that we used in order to calculate group statistics.

#### Bayesian Model Selection and Family Inference

For model selection, we estimated pointwise out-of-sample prediction accuracy for all fitted models separately for each participant by approximating leave-one-out cross-validation (LOO; corresponding to leave-one-trial-out per subject)^42, 84^. To do so, we applied Pareto smoothed importance sampling (PSIS) using the log-likelihood calculated from the posterior simulations of the parameter values as implemented by Vehtari et al.^42^. Sum PSIS-LOO scores for each model as well as information about *k̂* values – the estimated shape parameters of the generalized Pareto distribution – indicating the reliability of the PSIS-LOO estimate are depicted in **Supplementary Table S1**. As summarized in **Supplementary Table S1**, very few trials resulted in insufficient parameter values for *k̂* and thus potentially unreliable PSIS-LOO scores (on average 1.1 trials per subject with *k̂*>0.7 for the winning model)^42^. BMS on PSIS-LOO scores was performed on the group level, accounting for group heterogeneity in the model that best describes learning behavior^85^. This procedure provides the protected exceedance probability for each model (*pxp*), indicating how likely a given model is to have a higher probability of explaining the data than all other models in the comparison set. The Bayesian omnibus risk (*BOR*) indicates the posterior probability that model frequencies for all models are all equal to each other^85^. We also provide difference scores of PSIS-LOO in contrast to the model that won the BMS, which can be interpreted as a simple ‘fixed-effect’ model comparison (see **Supplementary Table S1)**^42, 84^. Model comparisons according to PSIS-LOO difference scores were qualitatively comparable to the BMS analyses for our data.

#### Posterior Predictive Checks and Statistical Analyses of Learning Parameters

First, posterior predictive checks were conducted by quantifying whether the predicted data could capture the variance in performance expectation ratings for each subject within each of the experimental conditions using regression analyses. Additionally, to assess whether the winning model captured the core effects in the behavioral data, we repeated the model-free analysis, which we had conducted on the behavioral data, with the data predicted by the winning model (see **Supplementary Results**).

Model parameters, i.e. learning rates, of the winning models for all experiments were analyzed on the group level. A repeated measures ANOVA was calculated on the learning rates with the factor Agent (Self [α_Self/PE+_, α_Self/PE-_] vs. Other [α_Other/PE+_, α_Other/PE-_]) and factor PE valence [pos| neg] (PE+ [α_Self/PE+_, α_Other/PE+_] vs. PE- [α_Self/PE-_, α_Other/PE-_]) as well as Group as a between-subject factor, testing whether learning about one’s own performance was more valence-specific than learning about the other person’s performance.

To associate learning biases with self-conscious affect, that is, embarrassment and pride, as well as self-esteem (SDQ-III subscale scores), we calculated a normalized learning rate valence bias score for self-related learning (Valence Learning Bias=(α_PE+(S)_ - α_PE-(S)_)/(α_PE+(S)_ + α_PE-(S)_))^5, 43, 44^. Spearman correlations were calculated between Valence Learning Bias, affect ratings, and self-esteem scores.

### Pupil Data Analysis

For the fMRI sample, eye-tracking data were assessed during scanning. Pupil diameter and gaze behavior were recorded non-invasively in one eye at 500 Hz using an MRI-compatible Eyelink-1000 plus device (SR Research, Kanata, ON, Canada) with manufacturer-recommended settings for calibration and blink detection. Due to insufficient pupillometry data quality, three participants had to be excluded from the analyses (final sample n=36). Pupil data were preprocessed by cutting out periods of blinks, and values in this gap were interpolated by piecewise cubic interpolation. The pupil trace was subsequently z-normalized over the whole session. To characterize the pupil dilation for each trial by a single value, we calculated a linear slope for each feedback phase of three seconds. Pupil traces were only analyzed for the Self condition, as onsets during feedback strongly differed between Agent conditions, rendering it impossible to make a meaningful comparison of pupil slopes between Agent conditions. Pupil slopes during self-related feedback phases for each trial were then entered into linear mixed models fitted by restricted maximum likelihood including PE valence [neg↗ pos] (continuous signed PE values) and PE surprise (continuous unsigned/ absolute PE values) as fixed effects and participant and PE valence [neg↗ pos] as random effects. Additionally, separate linear mixed models including embarrassment ratings, pride ratings or the Valence Learning Bias as well as their interaction with PE valence [neg↗ pos] were implemented to assess whether variance in individual pupil responses to positive and negative PEs (random PE valence [neg↗ pos] slopes) was explained by different emotional reactions and learning behavior (see Figure 2c).

### fMRI Data

#### fMRI Image Acquisition

Participants were scanned using a 3T Siemens MAGENTOM Skyra scanner (Siemens, München, Germany) at the Center of Brain, Behavior, and Metabolism (CBBM) at the University of Lübeck, Germany with 60 near-axial slices. An echo planar imaging (EPI) sequence was used for the acquisition of on average 1520 functional volumes (min=1395, max=1672) during each of the two sessions of the experiment, resulting in a total of on average 3040 functional volumes (TR=0.992s, TE=28ms, flip angle=60°, voxel size=3×3×3mm ^3^, simultaneous multi-slice factor 4). In addition, a high-resolution anatomical T1 image was acquired, which was used for normalization (voxel size=1×1×1mm^3^, 192×320×320mm^3^ field of view, TR=2.300s, TE=2.94ms, TI=900ms; flip angle=9°; GRAPPA factor 2; acquisition time 6.55 min).

#### FMRI data analyses

FMRI data were analyzed using SPM12 (www.fil.ion.ucl.ac.uk/spm). Field maps were reconstructed to obtain voxel displacement maps (VDMs). EPIs were corrected for timing differences of the slice acquisition, motion-corrected and unwarped using the corresponding VDMs to correct for geometric distortions and normalized using the forward deformation fields as obtained from the unified segmentation of the anatomical T1 image. The normalized volumes were resliced with a voxel size of 2×2×2 mm and smoothed with an 8 mm full-width-at-half-maximum isotropic Gaussian kernel. To remove low-frequency drifts, functional images were high-pass filtered at 1/384.

Statistical analyses were performed using a two-level, mixed-effects procedure. Three main GLMs were implemented on the first level. The first fixed-effects GLM included four epoch regressors modeling the hemodynamic responses to the different cue conditions (Ability: High vs. Low × Agent: Self vs. Other), weighted with the performance expectation ratings per trial as parametric modulator for each condition. Four regressors modeled the four feedback conditions (PE valence [pos| neg]: Positive vs. Negative × Agent: Self vs. Other), each weighted with the PE value for each trial. The estimation periods for Self and Other were modeled as two regressors, and emotion ratings phase and the instruction phase as separate regressors. To account for noise due to head movement, six additional regressors modeling head movement parameters were introduced and a constant term was included for each of the two sessions. The second first-level GLM differed only with respect to the feedback regressors. Here, only two regressors modeled feedback separately for Self and Other and two parametric modulators were included per condition, weighting feedback trials with PE valence [neg↗ pos] (continuous effect of the signed PE values) and PE surprise (continuous effect of the unsigned PE values). The third first-level GLM was set up to show that activation found in response to PEs was actually related to PEs and not only to the feedback level alone. Therefore, the parametric weights of the two feedback conditions in the second GLM were replaced by feedback level and performance expectation ratings, which allowed us to assess whether neural activity goes up with feedback level and down with performance expectation ratings, confirming a potential interpretation in terms of PE tracking^51^.

On the second level, for the first GLM model, beta images for the four feedback conditions were included in a flexible factorial design with two repeated measurement factors (PE valence [pos| neg] and Agent). Beta images for the parametric weights of feedback were extracted from the second and third first-level model for Self and Other. Separate repeated measures ANOVAs and one sample t-tests (for baseline contrasts) were implemented for PE valence [neg↗ pos] and PE surprise as well as feedback level and performance expectation level. Additional second-level models for the PE valence [neg↗ pos] contrast included the Valence Learning Bias, embarrassment, and pride ratings as between-subject covariates and assessed differential tracking of PEs depending on biased learning and self-conscious affect. A self-related Pupil Dilation Bias (average slope for positive PEs - average slope for negative PEs; higher scores indicate stronger pupil dilation for positive PEs) was also included as covariate in another second-level model to assess whether the neural response to negative vs. positive PEs was associated with the pupil dilation response.

We additionally performed psychophysiological interaction (PPI) analyses on the first level, investigating whether functional connectivity of the dAI, which is commonly activated during feedback processing independent of agent and feedback valence (conjunction of baseline contrasts: feedback Self ˄ feedback Other), would differ depending on the PE valence [neg↗ pos]. PPI analyses were computed separately for Self and Other and the resulting contrast images for the PPI effects were aggregated on the second level using two-sample t-tests contrasting PPI effects for Self vs. Other. For each participant, we defined 6-mm radius spherical ROIs, centered at the nearest local maximum for the conjunction contrast feedback Self ˄ feedback Other and located within 10 mm of the group maximum within the dAI, separately for the left dAI (x, y, z: −33 20 −4) and right dAI (x, y, z: 36 20 −7). By computing the first eigenvariate for all voxels within these ROIs that showed a positive effect for the conjunction (p<.500), we extracted the time course of activations and constructed PPI terms using the contrast for the parametric weights of PE valence [neg↗ pos] for Self or Other, respectively, resulting in four distinct PPI first-level GLMs. One participant was excluded from the PPI analyses for the right dAI, because no voxels survived the predefined threshold for eigenvariate extraction. The PPI term, along with the activation time course from the (left or right) dAI was included in a new GLM for each participant that also included all the regressors in the initial first-level GLM (four regressors for the different cue conditions, each weighted with the expected performance ratings; two feedback regressors for Self and Other with each two parametric modulators for PE valence [neg↗ pos] and PE surprise; two regressors for the estimation periods for Self and Other; one regressor for the emotion ratings phase; one regressor for the instruction phase; six regressors modeling head movement parameters; a constant term for each session). On the second level, we assessed whether there was a stronger functional coupling of the dAI with our predefined ROIs (Amygdala, mPFC, VTA/ SN) for the Self in contrast to the Other when PE valence [neg↗ pos] was more negative. Functional connectivity dynamics were also associated with learning behavior by calculating Spearman correlations for the Valence Learning Bias and the parameter estimates for the PPI effect of Self > Other derived from a sphere of 6mm around the peak voxels within our predefined ROIs.

#### Thresholding procedure and regions of interest

According to its suggested role as an integrative hub for motivated cognition and emotional behavior, the AI was defined as one of the regions of interest (ROIs)^38, 39^. Due to their specific functional associations, a bilateral ventral and a bilateral dorsal AI ROI was defined according to the three-cluster solution of Kelly and colleagues^86^. The bilateral amygdala was defined as another ROI and derived from the AAL atlas definition in the WFU PickAtlas^87^ due to its similar role for the attention-emotion interaction^38, 59^. The mPFC ROI was also derived from the AAL atlas in the WFU PickAtlas (label: bilateral frontal superior medial) due to its specific role during social learning and for biases in self-related belief updating reported in previous studies^29, 88^. Additionally, a VTA/ SN ROI, dopaminergic nuclei in the midbrain, was included^89, 90^, as dopamine signals motivationally important events, e.g. during reward learning^41^, and has been associated with biases in memory towards events that are of motivational significance^40^.

FMRI results were family-wise-error (FWE) corrected on the whole brain level unless otherwise mentioned, and all coordinates are reported in MNI space. As our predefined ROIs were chosen with respect to their involvement with the emotion-cognition link, we tested the effects of our covariates on PE valence [neg↗ pos] tracking and PPI effects within the ROIs. Anatomical labels of all resulting clusters were derived from the Automated Labeling Atlas Version 3.0^91^.

## Acknowledgments

We would like to thank Prof. Christoph W Korn for his very helpful comments and discussions on the manuscript. We are also grateful to Clara Gunzelmann and Rebecca Rocksien for their help with data collection. The research was funded by the German Research Foundation (Temporary Positions for Principal Investigators: MU 4373/1-1; Sachbeihilfe KR 3803/11-1) and the Medical Department of the University of Lübeck (J21-2018).

## Author contributions

L.M.P., N.C., F.M.P. and S.K., designed the research. L.M.P. and N.C. acquired the data. L.M.P. analyzed the data and prepared the manuscript. L.M.P., N.C., A.V.M., A.S., D.S., F.M.P., and S.K. discussed the data analyses and interpretation of the results and reviewed and edited the manuscript.

## Data availability

The data that support the findings of this study are available from the corresponding author upon reasonable request.

## Code availability

Code used to generate the analyses are available from the corresponding author upon reasonable request.

