## Supplementary Information for "Neurocomputational mechanisms of affected beliefs"

**Affiliations:**

**Supplementary Figures**

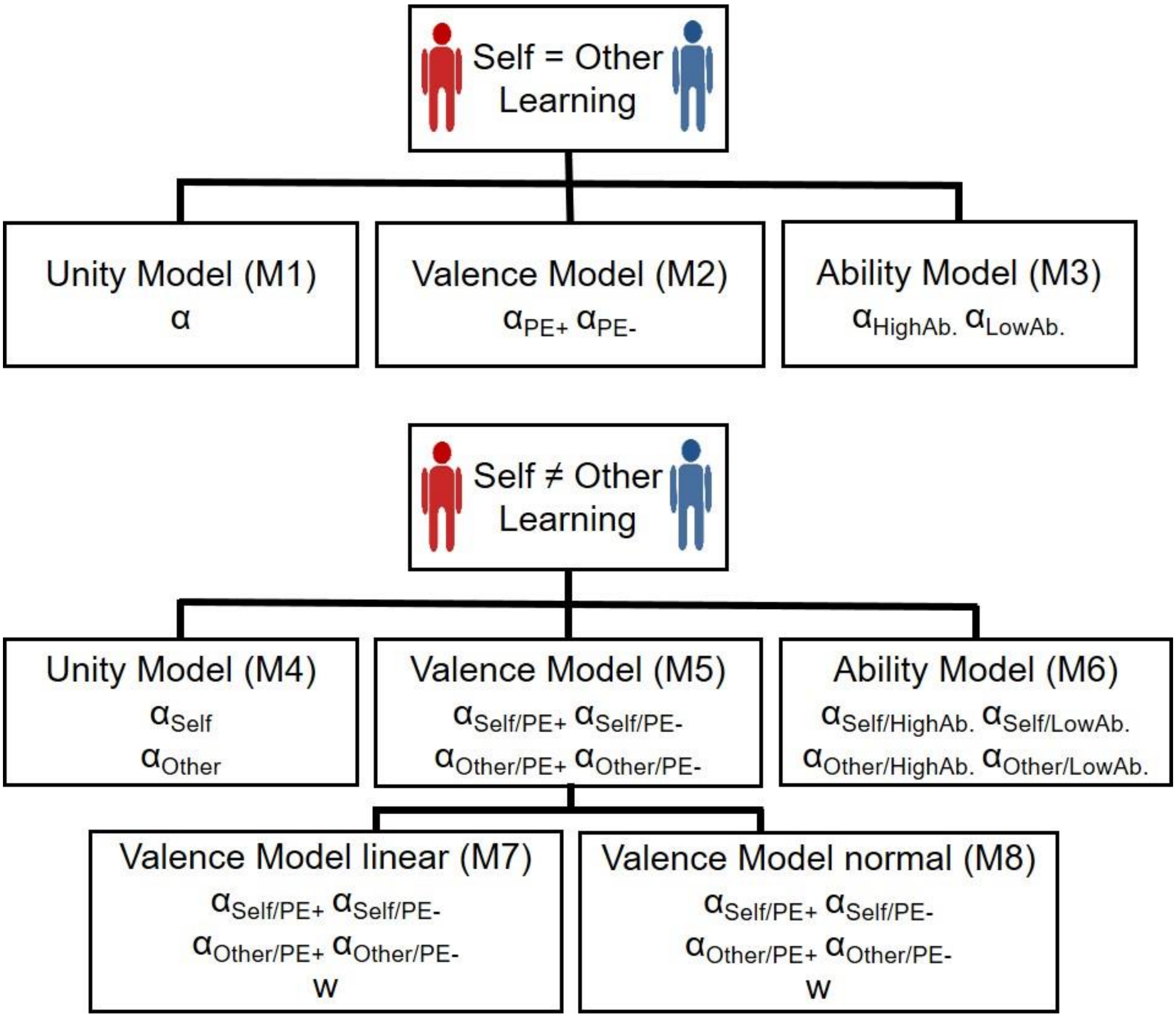


**Figure S1. Structure of the model space.** Two factors were distinguished that impact learning rates (α): the agent (self vs other) and the impact (no impact: Unity Model) of prediction error valence (Valence Model) or the ability condition (Ability Model). The Valence Model, winning model in previous studies (Müller-Pinzler et al., 2019), was extended by a decay factor (w) for the learning rates towards the ends of the feedback scale with a linear decrease (Valence Model linear) or a decrease following the relative probability density of the normal distribution (Valence Model normal; for more details see **Methods** and **Supplementary Methods** section).

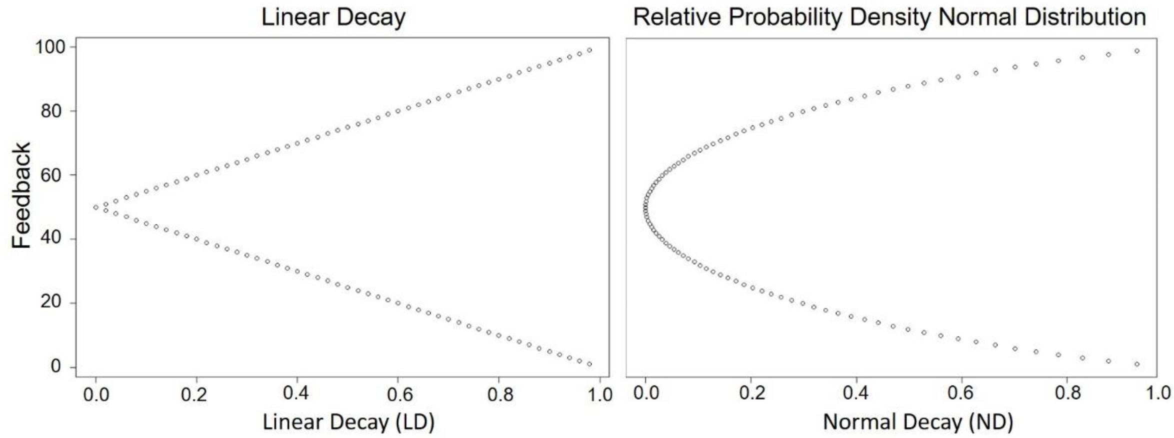


**Figure S2. Depiction of the linear decay (left) and the decay following the relative probability density of the normal distribution (right) for the different feedback values.** The values depicted here were introduced in the learning models and weighted by a weighting factor as described in the **Supplementary Methods** section.

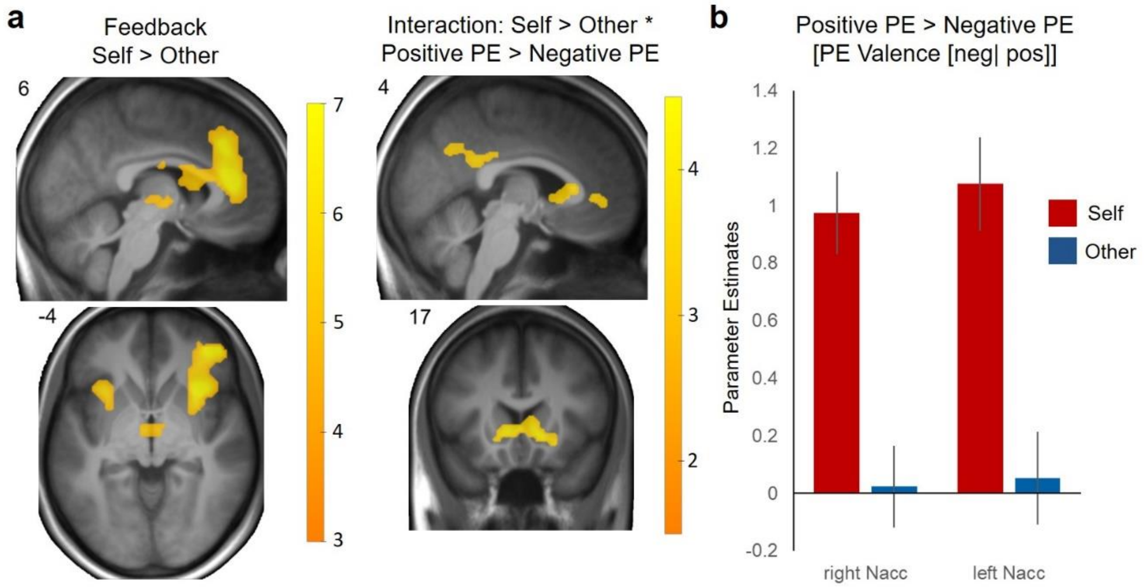


**Figure S3. Neural activations associated with feedback processing. a)** Self-related feedback vs. other-related feedback (left) was associated with an increased activation of the mPFC/ ACC, bilateral anterior insula and thalamus, among other regions ($p$ < .05, FWE-corrected). The interaction of Agent and PE valence [pos| neg] (([Self positive PE > Self negative PE] > [Other positive PE > Other negative PE]; right) resulted in activation of the angular gyrus, the bilateral NAcc/VS, the precuneus/ posterior cingulate cortex, and precentral gyrus (cluster-wise FWE-corrected with p < .05 at a cluster forming threshold of p < .001). **b)** Parameter estimates for the differences between feedback for positive vs. negative PEs derived from the peak voxels of the interaction effect depict the interaction in the left NAcc/ VS [x, y, z: -9 20 -1] and right NAcc/ VS [x, y, z: 12 20 -1]. Self-related feedback resulted in a valence-specific activation while other-related feedback did not.

### Supplementary Methods

#### Further information on the model space.

For the learning models the following PE delta-rule update equation (adapted Rescorla-Wagner model; Rescorla and Wagner, 1972) was used (EXP: Performance expectation rating, FB = feedback, PE = prediction error, α = learning rate):

$$EXP_{t+1} = EXP_t + \alpha\ PE_t;\ \text{while}\ PE_t = FB_t - EXP_t$$

Depending on the model (see **Figure S1** and **Methods** section) learning rates were adapted with respect to the different conditions (Agent, Ability condition or PE valence) and the initial beliefs about the own and the other participant's performance ($EXP_1$) were estimated as free parameters separately for Self and Other as well as both Ability conditions, resulting in four additional model parameters. The linear (LD) and normal decay (ND; values depicted in **Figure S2**) weighted by the weighting factor *w* that reduce the learning rates towards the ends of the scale were introduced in the learning models in the following way:

$$EXP_{t+1} = EXP_t + \alpha\ PE_t\,(1 - w\ LD);\ \text{for the linear decrease;}$$

$$EXP_{t+1} = EXP_t + \alpha\ PE_t\,(1 - w\ ND);\ \text{for the normal decrease.}$$

### Supplementary Results

#### Model free behavioral analyses reveal more negative self-evaluation.

First, we performed a model-free analysis to capture the basic effects observed in our behavioral data. Analyses of behavioral data and learning rates are based on the combined fMRI ($n$=39) and behavioral sample ($n$=30; total sample $N$=69). The Trial x Ability condition x Agent condition x Group ANOVA revealed a significant main effect of Ability condition ($F_{(1,67)}$=175.51, $p$<.001) and interaction of Trial x Ability condition ($F_{(19,1273)}$=108.87, $p$<.001), indicating that participants adapted their performance expectation ratings over time according to the feedback provided in each Ability condition (see **Figure 1c**). Moreover, there was a significant main effect of Agent condition ($F_{(1,67)}$=44.70, $p$=.001), indicating that participants evaluated their own performance more negatively than the other's performance. No significant interaction of Agent condition x Ability condition emerged ($F_{(1,67)}$=0.67, $p$=.415). The three-way interaction of Trial x Agent condition x Ability condition ($F_{(19,1273)}$=1.60, $p$=.047) revealed a significant effect, hinting at differential learning patterns between the Ability conditions for Self vs. Other. Finally, we found a significant main effect of Group ($F_{(1,67)}$=4.32, $p$=.041), indicating slightly higher ratings in the fMRI sample, but Group did not interact with any of the above-reported effects ($p$>.174).

#### Posterior predictive checks: Behavioral analyses on the predicted data.

To assess whether our winning model captured the core effects in our model free analysis, we let the parametrized winning model predict the time course of EXP for each participant, and compared these model predictions against the actual data (see **Figure 1c**). **Figure 1c** visually confirms the ability of the model to capture the observed data despite its small number of parameters. We repeated the behavioral analyses we had done on the actual behavioral data on the predicted data. The Trial x Ability condition x Agent condition x Group ANOVA on the predicted data revealed a significant main effect of Ability condition ($F_{(1,67)}$=205.28, $p$<.001) and interaction of Trial x Ability condition ($F_{(19,1273)}$=268.90, $p$<.001) replicating the effect that participants learned over time. More negative performance expectations for the self could also be replicated as indicated by the main effect of Agent ($F_{(1,67)}$=44.48, $p$<.001), while there was again no significant interaction of Agent condition x Ability condition ($F_{(1,67)}$=0.63, $p$=.429). The three-way interaction of Trial x Agent condition x Ability condition ($F_{(19,1273)}$=2.28, $p$=.001) showed a significant effect indicating differential learning patterns between the Ability conditions for self vs other. There was a significant main effect of Group ($F_{(1,67)}$=4.33, $p$=.041), but there was no interaction of Group with any of the effects reported above ($p$>.155). Repeating the behavioral analysis done on the model free data onto the predictions thus confirmed that it recapitulates the main effects in our data.

**Neural activations associated with feedback processing indicate a specific role of feedback valence during self-related learning.**

To examine the brain processes that underlie how people form self- and other-related ability beliefs, we compared neural activation during feedback processing as measured with fMRI. We found that the bilateral insula, anterior cingulate cortex, and thalamus (amongst others, see **Figure 3a** and **Supplementary Table S2**) were activated significantly more strongly for self-related compared to other-related performance feedback (i.e. Agent effect). This finding of heightened activity in brain regions that have been linked to arousal, but also to self-agency, potentially reflects a difference in the subjective salience of self- vs. other-related information (Craig, 2009; Späti et al., 2014; Sperduti et al., 2011). Compared to feedback for the Self, feedback for the Other resulted in stronger activation of the left and right middle temporal gyrus and precuneus/ middle cingulate gyrus (**Supplementary Table S2**).

Second, we compared self-related positive vs. negative feedback in order to examine how the valence of information affected neural processing (categorical PE valence [pos| neg] effect). We found significantly stronger activations of the left and right nucleus accumbens/ ventral striatum (NAcc/VS), bilateral angular gyrus, medial prefrontal cortex (mPFC), and precuneus/ posterior cingulate cortex (PCC) for positive PE valence than for negative PE valence [pos| neg] (see **Supplementary Table S2**). This valence effect was unique for the processing of self-related information and did not emerge for other-related performance feedback (no significant clusters for the PE valence [pos| neg] effect for Other; $p$ < .001). The opposite contrast, negative vs. positive PE valence [pos| neg], yielded no significant activations, either for self-related or for other-related information. When testing the interaction of Agent x PE valence [pos| neg], we found increased activation for self-related positive vs. negative feedback ([Self positive PE > Self negative PE] > [Other positive PE > Other negative PE]) in the angular gyrus (see **Supplementary Table S2**), and at a more lenient threshold also the bilateral NAcc/VS, the precuneus/ PCC, and precentral gyrus (cluster-wise FWE-corrected with $p$ < .05 at a cluster forming threshold of $p$ < .001; see **Figure 3a/ b** and **Supplementary Table S3**).

**Specific associations of embarrassment and pride with neural activity in response to self-related prediction error valence.**

To test whether embarrassment and pride had independent effects on neural activity in response to self-related PE valence within our predefined ROIs, we extracted parameter estimated for the effect of the parametric weights for PE valence [neg↗pos] for each whole ROI. Mean parameter estimates for the whole ROIs were then entered into regression models predicting the neural activity with both affect ratings simultaneously. We found independent effects of pride ($\beta$=0.36, $t_{(36)}$=2.63, $p$=.012) and embarrassment ($\beta$=-0.39, $t_{(36)}$=-2.82, $p$=.008; $R^2$=.33, $F_{(2,36)}$=8.94, $p$<.001) within the amygdala. We also found independent effects of pride ($\beta$=0.43, $t_{(36)}$=3.17, $p$=.003) and embarrassment ($\beta$=-0.36, $t_{(36)}$=-2.63, $p$=.013; $R^2$=.36, $F_{(2,36)}$=10.10, $p$<.001) within the dAI. For the vAI we found a significant effect of pride ($\beta$=0.38, $t_{(36)}$=2.61, $p$=.013) and a trend-wise effect of embarrassment ($\beta$=-0.29, $t_{(36)}$=-2.01, $p$=.052; $R^2$=.26, $F_{(2,36)}$=6.47, $p$=.004). Independent effects for pride ($\beta$=0.39, $t_{(36)}$=2.85, $p$=.007) and embarrassment ($\beta$=-0.40, $t_{(36)}$=-2.91, $p$=.006; $R^2$=.36, $F_{(2,36)}$=9.97, $p$<.001) were also present for the mPFC and we also found independent effects of pride ($\beta$=0.32, $t_{(36)}$=2.39, $p$=.022) and embarrassment ($\beta$=-0.46, $t_{(36)}$=-3.37, $p$=.002; $R^2$=.36, $F_{(2,36)}$=10.20, $p$<.001) for the VTA/ SN.

### Supplementary Tables

*Table S1.* PSIS-LOO Scores

| Model | PSIS-LOO | LOO-SE | LOO-Diff (SE-Diff) | % of $\hat{k} > 0.7$ | No. Est. Parameters |
|---|---|---|---|---|---|
| Mean Model (M0) | -2644.4 | 319.7 | 1436.1 (142.8) | 0.07 | 4 |
| **Self = Other** | | | | | |
| Unity Model (M1) | -1801.3 | 396.5 | 593.0 (109.0) | 0.47 | 5 |
| Context Model (M2) | -1681.2 | 367.8 | 472.9 (80.6) | 0.58 | 6 |
| Valence Model (M3) | -1679.3 | 388.0 | 470.9 (93.9) | 0.74 | 6 |
| **Self ≠ Other** | | | | | |
| Unity Model (M4) | -1621.2 | 363.6 | 412.9 (75.5) | 0.34 | 6 |
| Context Model (M5) | -1599.9 | 372.6 | 391.6 (69.2) | 1.43 | 8 |
| Valence Model (M6) | -1346.4 | 333.6 | 138.1 (39.0) | 0.53 | 8 |
| ext. Valence Model (M7) | -1251.4 | 349.2 | 43.1 (16.7) | 1.58 | 9 |
| ext. Valence Model (M8) | -1208.3 | 357.7 | - | 1.39 | 9 |

*Note*. LOO = sum PSIS-LOO, approximate leave-one-out cross-validation (LOO) using Pareto-smoothed importance sampling (PSIS); LOO-SE = Standard error of PSIS-LOO; LOO-Diff (SE-Diff) = Difference in expected predictive accuracy (PSIS-LOO) for all models from the model with the highest PSIS-LOO (extended Valence Model M8) and standard errors of differences; percentage of $\hat{k}$ - estimated shape parameters of the generalized Pareto distribution - exceeding 0.7 (all according to Vehtari et al., 2016); No. Est. Parameters = number of estimated parameters in the model.

*Table S2.* Activations Associated with Feedback Processing

| Contrasts/ Brain Regions | Side | Cluster Size | MNI Coordinates | | | *T* | *p* |
|---|---|---|---|---|---|---|---|
| | | | x | y | z | | |
| **Self > Other** | | | | | | | |
| Anterior Cingulate Gyrus/ Paracingulate Gyrus | R/L | 1626 | 6 | 38 | 14 | 8.58 | <.001 |
| Frontal Pole | R | | 45 | 47 | -7 | 8.33 | <.001 |
| Insular Cortex / Frontal Orbital Cortex | R | | 36 | 17 | -7 | 7.81 | <.001 |
| Insular Cortex / Frontal Orbital Cortex | L | 126 | -36 | 17 | -10 | 7.14 | <.001 |
| Anterior/ Posterior Supramarginal Gyrus | R | 234 | 48 | -37 | 47 | 6.73 | <.001 |
| Inferior Frontal Gyrus, pars opercularis/ Precentral Gyrus | R | 74 | 51 | 14 | 14 | 5.8 | .001 |
| Thalamus | R/L | 54 | 6 | -7 | -1 | 5.23 | .005 |
| | | | -3 | -10 | -4 | 5.21 | .005 |
| | | | 3 | -22 | -1 | 5.03 | .011 |
| **Other > Self** | | | | | | | |
| Anterior/ Posterior Middle Temporal Gyrus | L | 104 | -60 | -10 | -13 | 6.85 | <.001 |
| Angular Gyrus / Superior Lateral Occipital Gyrus | R | 100 | 57 | -58 | 23 | 6.68 | <.001 |
| Precuneous Cortex / Posterior Cingulate Cortex | R/L | 176 | 3 | -52 | 35 | 6.54 | <.001 |
| Posterior Cingulate Cortex/ Precuneous Cortex | | | -12 | -46 | 35 | 4.83 | .022 |
| Anterior Middle/ Anterior Superior Temporal Gyrus | R | 69 | 60 | -1 | -19 | 6.41 | <.001 |
| Temporal Pole | | | 51 | 14 | -31 | 5.98 | <.001 |
| **Self: Positive PE > Negative PE** | | | | | | | |
| Angular Gyrus/ Superior Parietal Lobule | L | 612 | -42 | -55 | 44 | 8.01 | <.001 |
| Superior Parietal Lobule/Superior Lateral Occipital Cortex | | | -36 | -58 | 56 | 7.32 | <.001 |
| Angular Gyrus/ Superior Lateral Occipital Cortex | R | 377 | 48 | -58 | 38 | 7.51 | <.001 |
| Angular Gyrus/ Superior Parietal Lobule | | | 45 | -49 | 53 | 5.81 | .001 |
| Anterior/ Posterior Supramarginal Gyrus | | | 51 | -37 | 50 | 5.69 | .001 |
| Superior Frontal Gyrus/ Middle Frontal Gyrus | L | 474 | -15 | 29 | 53 | 7.45 | <.001 |
| Middle Frontal Gyrus/ Superior Frontal Gyrus | | | -36 | 20 | 50 | 6.43 | <.001 |
| Middle Frontal Gyrus | | | -39 | 26 | 41 | 6.04 | <.001 |
| Caudate | R | 434 | 12 | 20 | 2 | 6.83 | <.001 |
| Caudate / Accumbens | L | | -9 | 20 | 2 | 6.83 | <.001 |
| Paracingualte Gyrus/ Anterior Cigulate Gyrus | R/L | | 0 | 47 | -1 | 5.81 | .001 |
| Posterior/ Anterior Cingulate Gyrus | R/L | 436 | 0 | -28 | 32 | 6.14 | <.001 |
| Posterior Cingulate Gyrus/ Precuneous Cortex | | | -6 | -55 | 29 | 6.03 | <.001 |
| Posterior Cingulate Gyrus | | | -3 | -37 | 29 | 5.61 | .001 |
| **Interaction: Self > Other, Positive PE > Negative PE** | | | | | | | |
| Angular Gyrus/ Superior Lateral Occipital Cortex | R | 19 | 48 | -58 | 41 | 5.28 | .004 |
| Angular Gyrus/ Superior Parietal Lobule | L | 9 | -42 | -55 | 44 | 4.78 | .027 |
| | | | -42 | -55 | 53 | 4.76 | .028 |
| Putamen/ Pallidum | R | 1 | 18 | 5 | -10 | 4.67 | .038 |

*Note*. Cluster extends refer to $p < .05$, FWE corrected for the whole brain and *p*-values are FWE for the whole brain, respectively. Only clusters with more than 50 voxels are reported. For the interaction contrast ([Self Positive PE > Self Negative PE] > [Other Positive PE > Other Negative PE]) all clusters are reported.

*Table S3.* Activations Associated with Feedback Processing: Interaction of Agent * PE Valence

| Contrasts/ Brain Regions | Side | Cluster Size | MNI Coordinates | | | *T* | *p* |
|---|---|---|---|---|---|---|---|
| | | | x | y | z | | |
| **Interaction: Self > Other, Positive > Negative** | | | | | | | |
| Angular Gyrus/ Superior Lateral Occipital Cortex | R | 229 | 48 | -58 | 41 | 5.28 | .002 |
| | | | 57 | -61 | 20 | 3.55 | |
| Angular Gyrus / Superior Parietal Lobule | | | | | | | |
| | L | 303 | -42 | -55 | 44 | 4.78 | .001 |
| | | | -42 | -55 | 53 | 4.76 | |
| Angular Gurus / Posterior Supramarginal Gyrus | | | -48 | -55 | 29 | 4.57 | |
| Putamen/ Pallidum | R | 380 | 18 | 5 | -10 | 4.67 | <.001 |
| Caudate / Accumbens | R | | 12 | 20 | -1 | 4.56 | |
| Caudate / Accumbens | L | | -9 | 20 | -1 | 4.55 | |
| Precentral gyrus | L | 162 | -18 | -19 | 53 | 3.95 | .008 |
| | | | -27 | -13 | 44 | 3.79 | |
| | | | -24 | -25 | 41 | 3.71 | |
| Posterior Cingulate Gyrus/ Precuneous Cortex | R/L | 296 | -15 | -43 | 32 | 3.91 | .001 |
| Precuneous Cortex/ Posterior Cingulate Gyrus | | | -3 | -58 | 35 | 3.87 | |
| Posterior Cingulate Gyrus/ Precuneous Cortex | | | 6 | -43 | 26 | 3.73 | |
| Cerebellum Left Crus I / Crus II | L | 154 | -12 | -82 | -25 | 3.73 | .009 |
| Occipital Fusiform Gyrus / Cerebellum Left Crus I | | | -30 | -79 | -1 | 3.69 | |
| Occipital Fusiform Gyrus / Inferior Lateral Occipital Cortex | | | -30 | -85 | -10 | 3.65 | |

*Note*. Cluster extends refer to $p < .001$, uncorrected and *p*-values are FWE corrected on the cluster level.

*Table S4.* Activations Associated with the Parametric Weights of PE Surprise

| Contrasts/ Brain Regions | Side | Cluster Size | MNI Coordinates | | | *T* | *p* |
|---|---|---|---|---|---|---|---|
| | | | x | y | z | | |
| **Self: PE Surprise** | | | | | | | |
| Paracingulate Gyrus/ Superior Frontal Gyrus | L /R | 237 | -6 | 50 | 20 | 5.75 | .002 |
| Temporal Pole/ Superior Frontal Gyrus | | | 3 | 56 | 14 | 4.39 | |
| Temporal Pole/ Frontal Orbital Cortex | | | | | | | |
| | L | 107 | -39 | 17 | -22 | 5.57 | .028 |
| | | | -30 | 11 | -28 | 5.10 | |
| Frontal Orbital Cortex / Insular Cortex | R | 165 | 30 | 23 | -13 | 4.35 | .009 |
| Temporal Pole | | | 51 | 14 | -28 | 4.29 | |
| Frontal Orbital Cortex | | | 39 | 26 | -16 | 4.23 | |
| **Other: PE Surprise** | | | | | | | |
| Temporal Pole/ Frontal Orbital Cortex | L | 220 | -33 | 17 | -28 | 6.90 | .001 |
| Anterior/Posterior Middle Temporal Gyrus | | | -63 | -10 | -19 | 5.17 | |
| Temporal Pole/ Anterior Middle Temporal Cortex | | | -57 | 2 | -25 | 4.56 | |
| Temporal Pole/ Frontal Orbital Cortex | R | 279 | 39 | 20 | -28 | 6.75 | <.001 |
| Temporal Pole | | | 48 | 17 | -31 | 5.27 | |
| Temporal Pole/ Anterior Middle Temporal Cortex | | | 51 | 8 | -31 | 4.86 | |
| Angular Gyrus/ Posterior Supramarginal Gyrus | R | 291 | 48 | -46 | 26 | 6.48 | <.001 |
| Superior Lateral Occipital Cortex | | | 39 | -79 | 26 | 4.14 | |
| Superior/ Inferior Lateral Occipital Cortex | | | 51 | -67 | 23 | 3.71 | |
| Superior Frontal Gyrus/ Frontal Pole | R/L | 510 | 6 | 53 | 26 | 6.41 | <.001 |
| Superior Frontal Gyrus/ Paracingulate Gyrus | | | -3 | 53 | 23 | 6.24 | |
| Frontal Pole/ Superior Frontal Gyrus | | | 9 | 47 | 47 | 5.50 | |
| Posterior Supramarginal Gyrus/ Angular Gyrus | L | 237 | -51 | -49 | 17 | 5.35 | .001 |
| Angular Gyrus/ Superior Lateral Occipital Gyrus | | | -57 | -58 | 29 | 4.96 | |
| Posterior Supramarginal Gyrus/ Angular Gyrus | | | -60 | -49 | 32 | 4.52 | |

*Note*. PE surprise refers to the unsigned prediction error values as parametric modulator for the feedback phase. Cluster extends refer to $p < .001$, uncorrected and $p$-values are FWE corrected on the cluster level.

*Table S5.* Activations Associated with the Parametric Weights of PE Valence

| Contrasts/ Brain Regions | Side | Cluster Size | MNI Coordinates | | | *T* | *p* |
|---|---|---|---|---|---|---|---|
| | | | x | y | z | | |
| **Self: PE Valence** | | | | | | | |
| Superior/ Middle Frontal Gyrus | L | 197 | -15 | 29 | 53 | 7.07 | <.001 |
| Middle/ Superior Frontal Gyrus | | | -36 | 17 | 50 | 6.38 | .002 |
| Middle Frontal Gyrus | | | -39 | 23 | 38 | 5.75 | .009 |
| Superior Parietal Lobule/Superior Lateral Occipital Cortex | L | 199 | -36 | -58 | 56 | 6.96 | <.001 |
| Angular Gyrus/ Posterior Supramarginal Gyrus | | | -45 | -55 | 35 | 6.77 | .001 |
| Caudate / Accumbens | L | 139 | -9 | 20 | -1 | 6.76 | .001 |
| Caudate / Accumbens | R | | 12 | 17 | -1 | 6.41 | .002 |
| Superior Lateral Occipital Gyrus/ Angular Gyrus | R | 121 | 48 | -61 | 41 | 6.48 | .001 |
| Posterior Supramarginal Gyrus/ Angular Gyrus | | | 51 | -46 | 47 | 6.34 | .002 |
| Superior Parietal Lobule/ Angular Gyrus | | | 39 | -55 | 56 | 5.44 | .020 |
| Postcentral Gyrus/ Superior Parietal Lobule | L | 50 | -45 | -34 | 56 | 6.08 | .004 |
| Postcentral Gyrus/ Posterior Supragarginal Gyrus | | | -45 | -28 | 41 | 5.53 | .016 |
| **Self > Other: PE Valence** | | | | | | | |
| Accumbens | L | 19 | -9 | 26 | -1 | 5.77 | 0.008 |
| Caudate/ Accumbens | R | 2 | 12 | 17 | -4 | 5.23 | 0.034 |

*Note*. PE valence refers to the signed prediction error values as parametric modulator for the feedback phase. Cluster extends refer to $p < .05$, FWE corrected for the whole brain and *p*-values are FWE corrected for the whole brain, respectively. Only clusters with more than 50 voxels are reported for the Self: PE Valence contrast.

*Table S6.* Activations Associated with the Parametric Weights of Feedback and Self-Related Expectations

| Contrasts/ Brain Regions | Side | Cluster Size | MNI Coordinates | | | *T* | *p* |
|---|---|---|---|---|---|---|---|
| | | | x | y | z | | |
| **Self: Feedback Value** | | | | | | | |
| Caudate/ Accumbens | L | 252 | -6 | 17 | 2 | 7.63 | <.001 |
| Caudate/ Accumbens | R | | 12 | 17 | -1 | 7.21 | <.001 |
| Superior Frontal Gyrus | L | 312 | -15 | 29 | 56 | 7.40 | <.001 |
| Middle Frontal Gyrus/ Superior Frontal Gyrus | | | -36 | 14 | 50 | 7.34 | <.001 |
| Middle Frontal Gyrus | | | -39 | 23 | 38 | 6.79 | .001 |
| Angular Gyrus/ Posterior Supramarginal Gyrus | L | 314 | -45 | -55 | 32 | 6.75 | .001 |
| | | | -45 | -55 | 41 | 6.61 | .001 |
| Superior Parietal Lobule/ Superior Lateral Occipital Cortex | | | -36 | -58 | 56 | 6.52 | .001 |
| Angular Gyrus/ Superior Lateral Occipital Cortex | R | 220 | 45 | -58 | 38 | 6.74 | .001 |
| Posterior Supramarginal Gyrus/ Angular Gyrus | | | 54 | -43 | 47 | 6.65 | .001 |
| | | | 48 | -43 | 38 | 5.88 | .007 |
| Middle Frontal Gyrus | R | 101 | 39 | 23 | 41 | 5.90 | .001 |
| Middle Frontal Gyrus/ Superior Frontal Gyrus | | | 33 | 23 | 50 | 6.51 | .007 |
| Superior Frontal Gyrus/ Middle Frontal Gyrus | | | 27 | 23 | 56 | 5.77 | .009 |
| | R/L | 49 | -3 | 53 | -1 | 6.10 | .004 |
| Paracingulate Gyrus/ Frontal Medial Cortex | | | -3 | 38 | 5 | 5.62 | .014 |
| Cerebellum Crus I/ Crus II | R | 49 | 39 | -64 | -40 | 6.00 | .005 |
| | | | 45 | -73 | -37 | 5.77 | .009 |
| | | | 36 | -73 | -46 | 5.17 | .045 |
| Postcentral Gyrus/ Superior Parietal Lobule | L | 64 | -45 | -34 | 56 | 5.98 | .005 |
| Postcentral Gyrus/ Anterior Supramarginal Gyrus | | | -45 | -28 | 41 | 5.53 | .018 |
| Anterior Supramarginal Gyrus/ Postcentral Gyrus | | | -48 | -34 | 47 | 5.42 | .023 |
| **Self: Expectation (negative associations)** | | | | | | | |
| Middle Frontal Gyrus | L | 27 | -36 | 14 | 53 | 6.73 | .001 |
| Accumbens/ Caudate | L | 23 | -9 | 20 | -4 | 6.33 | .003 |
| Superior Frontal Gyrus/ Middle Frontal Gyrus | L | 48 | -18 | 26 | 56 | 6.21 | .004 |
| Superior Frontal Gyrus/ Frontal Pole | | | -6 | 38 | 50 | 5.72 | .016 |
| Angular Gyrus/ Superior Lateral Occipital Cortex | R | 20 | 45 | -58 | 38 | 6.09 | .006 |
| Accumbens/ Caudate | R | 22 | 12 | 17 | -7 | 6.04 | .007 |
| Cerebellum Crus I/ Crus II | R | 32 | 42 | -73 | -37 | 5.93 | .009 |
| Angular Gyrus/ Posterior Supramarginal Gyrus | L | 6 | -39 | -52 | 35 | 5.70 | .017 |

*Note.* Cluster extends refer to $p < .05$, FWE corrected for the whole brain and $p$-values are FWE for the whole brain, respectively. Only clusters with more than 50 voxels are reported for the Self: Feedback Value contrast.

193

*Table S7.* Covariates Associated with Individual Differences in Valence Specific Prediction Error Tracking

| Covariates/ Regions of Interest | Side | Cluster Size | MNI Coordinates | | | *T* | *p* |
|---|---|---|---|---|---|---|---|
| | | | x | y | z | | |
| **Valence Learning Bias** | | | | | | | |
| Amygdala | R | 10 | 30 | 2 | -28 | 4.05 | .007 |
| | L | 7 | -30 | 5 | -19 | 3.92 | .010 |
| Dorsal Anterior Insula | R | 19 | 39 | 20 | -7 | 3.67 | .007 |
| | L | 28 | -30 | 11 | -19 | 4.31 | .004 |
| Ventral Anterior Insula | R | 1 | 33 | 11 | -16 | 3.54 | .033 |
| | L | 5 | -33 | 11 | -16 | 4.29 | .005 |
| Ventral Tegmental Area/ Substantia Nigra | R/L | 13 | 6 | -13 | -19 | 4.20 | .009 |
| | | 2 | 6 | -25 | -16 | 3.70 | .028 |
| | | 1 | 15 | -13 | -19 | 3.58 | .037 |
| Medial Prefrontal Cortex | R/L | 6 | 12 | 50 | 17 | 4.36 | .018 |
| | | 9 | -9 | 50 | 20 | 4.19 | .027 |
| | | 1 | -15 | 50 | 8 | 3.93 | .050 |
| **Pupil Valence Bias** | | | | | | | |
| Amygdala | R | 20 | 30 | -1 | -25 | 3.90 | .012 |
| | L | 5 | -30 | -1 | -22 | 3.44 | .034 |
| Dorsal Anterior Insula | R | 1 | 42 | 14 | -13 | 3.37 | .043 |
| | L | 6 | -36 | 5 | -13 | 3.76 | .018 |
| | | 2 | -33 | 5 | -1 | 3.32 | .048 |
| Ventral Anterior Insula | R | 3 | 42 | -1 | -19 | 4.43 | .004 |
| | | 2 | 42 | 11 | -16 | 3.55 | .035 |
| | L | 3 | -36 | 5 | -16 | 3.83 | .018 |
| Ventral Tegmental Area/ Substantia Nigra | R/L | 3 | 15 | -13 | -19 | 3.69 | .032 |
| | | 1 | -12 | -31 | -16 | 3.50 | .049 |
| **Embarrassment** | | | | | | | |
| Amygdala | R | 1 | 30 | 2 | -28 | 3.34 | .040 |
| | L | 4 | -18 | -7 | -16 | 3.69 | .018 |
| | | 4 | -27 | 2 | -19 | 3.34 | .040 |
| Dorsal Anterior Insula | R | 8 | 39 | 14 | -10 | 3.56 | .027 |
| | L | | -33 | 17 | 2 | 3.21 | .059 |
| Ventral Anterior Insula | R | | 33 | 11 | -16 | 3.08 | .094 |
| | L | | -33 | 5 | -16 | 3.11 | .089 |
| Ventral Tegmental Area/ Substantia Nigra | R/L | 79 | 9 | -25 | -19 | 4.49 | .004 |
| | | 14 | -12 | -28 | -7 | 3.81 | .023 |
| | | 2 | -15 | -10 | -13 | 3.78 | .025 |
| Medial Prefrontal Cortex | R/L | | 9 | 59 | 20 | 3.79 | .071 |
| **Pride** | | | | | | | |
| Amygdala | R | 5 | 33 | 2 | -28 | 3.39 | .037 |

| | | | | | | | |
|---|---|---|---|---|---|---|---|
| | | 1 | 21 | 2 | -13 | 3.29 | .045 |
| | L | | -21 | -1 | -16 | 3.20 | .055 |
| Dorsal Anterior Insula | R | 29 | 39 | 17 | -7 | 3.68 | .020 |
| | | 1 | 33 | 14 | -13 | 3.29 | .050 |
| | L | 22 | -36 | 20 | 2 | 3.94 | .011 |
| Ventral Anterior Insula | R | 6 | 42 | -7 | -13 | 3.94 | .013 |
| | | 1 | 33 | 11 | -16 | 3.52 | .036 |
| | L | 1 | -42 | 11 | -7 | 3.54 | .035 |
| Ventral Tegmental Area/ Substantia Nigra | R/L | 10 | 0 | -19 | -19 | 3.95 | .017 |
| | | 1 | 15 | -16 | -19 | 3.52 | .044 |
| | | 1 | 21 | -16 | -10 | 3.47 | .049 |
| Medial Prefrontal Cortex | R/L | 6 | 15 | 53 | 14 | 4.17 | .030 |
| | | 6 | -9 | 50 | 23 | 4.02 | .042 |

*Note*. Cluster extends refer to $p < .05$, FWE corrected within ROIs and $p$-values are FWE corrected within ROIs, respectively. Trendwise effects are indicated in grey.

*Table S8.* Differential Functional Connectivity of the Dorsal Anterior Insula Associated with Prediction Error Valence

| Covariates/ Regions of interest | Side | Cluster Size | MNI Coordinates x | y | z | *T* | *p* |
|---|---|---|---|---|---|---|---|
| **PPI Right Dorsal Anterior Insula** | | | | | | | |
| Amygdala | R | 6 | 33 | -1 | -31 | 4.46 | .003 |
| | L | 2 | -27 | -4 | -25 | 3.89 | .013 |
| | | 5 | -24 | 2 | -13 | 3.68 | .022 |
| Ventral Tegmental Area/ Substantia Nigra | R/L | 5 | -18 | -16 | -13 | 4.07 | .015 |
| Medial Prefrontal Cortex | R/L | 11 | -9 | 35 | 53 | 4.95 | .005 |
| | | 5 | -6 | 59 | 26 | 4.62 | .012 |
| **PPI Left Dorsal Anterior Insula** | | | | | | | |
| Amygdala | L | 3 | -30 | -4 | -22 | 3.76 | .019 |
| Ventral Tegmental Area/ Substantia Nigra | R/L | 3 | -9 | -13 | -13 | 3.66 | .042 |

*Note*. Stronger functional connectvity for negative vs positive PEs for self- vs other-related feedback. Cluster extends refer to $p < .05$, FWE corrected within ROIs and $p$-values are FWE corrected within ROIs, respectively.

*Table S9.* Sample characteristics

| | fMRI Sample | | Behavioral Sample | | p |
|---|---|---|---|---|---|
| | Mean | SD | Mean | SD | |
| Age | 22.30 | 2.65 | 23.30 | 3.97 | .234 |
| Self-esteem | 6.24 | 0.84 | 6.07 | 1.26 | .522 |

*Note.* Sample characteristics for both samples. SD = standard devation; fMRI Sample: n=39, Behavioral Sample: n = 30; p-value refers to a two sample t-test, df = 67.
